# Distinct and common features of numerical and structural chromosomal instability across different cancer types

**DOI:** 10.1101/2021.10.15.464567

**Authors:** Xiaoxiao Zhang, Maik Kschischo

**Affiliations:** Department of Mathematics and Technology, University of Applied Sciences Koblenz, Joseph-Rovan-Allee 2, 53424 Remagen, Germany; Department of Informatics, Technical University of Munich, Munich, 81675, Germany

**Keywords:** whole chromosomal instability, structural chromosomal instability, whole genome doubling, integrative analysis, PI3K oncogenic activation

## Abstract

A large proportion of tumours is characterised by numerical or structural chromosomal instability (CIN), defined as an increased rate of gaining or losing whole chromosomes (W-CIN) or of accumulating structural aberrations (S-CIN). Both W-CIN and S-CIN are associated with tumourigenesis, cancer progression, treatment resistance and clinical outcome. Although W-CIN and S-CIN can co-occur, they are initiated by different molecular events. By analysing tumour genomic data from 33 cancer types, we show that the majority of tumours with high levels of W-CIN underwent whole genome doubling, whereas S-CIN levels are strongly associated with homologous recombination deficiency. Both CIN phenotypes are prognostic in several cancer types. Most drugs are less efficient in high-CIN cell lines, but we also report compounds and drugs which should be investigated as targets for W-CIN or S-CIN. By analysing associations between CIN and bio-molecular entities with pathway and gene expression levels, we complement gene signatures of CIN and report that the drug resistance gene *CKS1B* is strongly associated with S-CIN. Finally, we propose a potential copy number-dependent mechanism to activate the *PI3K* pathway in high-S-CIN tumours.

**Simple summary:** Many cancer cells are chromosomally unstable, a phenotype describing a tendency for accumulating chromosomal aberrations. Entire chromosomes tend to be gained or lost, which is called whole chromosome instability (W-CIN). Structural chromosomal instability (S-CIN) describes an increased rate of gaining, losing or translocating smaller parts of chromosomes. Here, we analyse data from 33 cancer types to find differences and commonalities between W-CIN and S-CIN. We find that W-CIN is strongly linked to whole genome doubling (WGD), whereas S-CIN is associated with a specific DNA damage repair pathway. Both W-CIN and S-CIN are difficult to target using currently available compounds and have distinct prognostic values. The activity of the drug resistance gene *CKS1B* is associated with S-CIN, which merits further investigation. In addition, we identify a potential copy number-based mechanism promoting signalling of the important *PI3K* cancer pathway in high-S-CIN tumours.

## 2. Introduction

A large proportion of human tumours exhibits abnormal karyotypes with gains and losses of whole chromosomes or structural aberrations of parts of chromosomes [1–3]. In many cases, these karyotypic changes are the result of ongoing chromosomal instability (CIN), which is defined as an increased rate of chromosomal changes. Accordingly, two major forms of CIN can be distinguished: Whole chromosome instability (W-CIN), which is also called numerical CIN, refers to the ongoing acquisition of gains and losses of whole chromosomes. Structural CIN (S-CIN) is characterised by an increased rate of acquiring structural changes in chromosomes including, amongst other things, amplifications and deletions, inversions, duplications and balanced or unbalanced translocations [1–4]. CIN is to be distinguished from polyploidy, where the whole set of chromosomes is increased. In cross-sectional tumour samples, W-CIN manifests itself by an abnormal and unequal number of chromosomes, whereas the S-CIN phenotype is characterised by segmental aneuploidy, i.e., gains and losses of chromosome segments. Although W-CIN can induce S-CIN and vice versa, both types of CIN arise through distinct molecular characteristics. Whilst W-CIN is caused by chromosome missegregation during mitosis, S-CIN is commonly attributed to errors in the repair of DNA double-strand breaks [5, 6]. Both types of CIN are intimately related to DNA replication stress [7, 8], which not only induces CIN [9, 10], but also occurs as an immediate short-term response to aneuploidy and CIN [11].

Aneuploidy and CIN have typically detrimental effects on cell fitness and proliferation [5, 11, 12]. Therefore, it was unclear why CIN is often associated with poor patient survival and more aggressive disease progression [1, 13, 14]. Stratification of breast cancer patient samples into low, intermediate and high CIN groups revealed that patients with intermediate levels of CIN had the worst survival, whereas the low and high CIN groups had a better prognosis [15, 16]. These results hinted at mechanisms for tolerating CIN in order to survive the stresses provoked by chromosomal aberrations. The CIN tolerance mechanisms are currently not completely understood [17], but one important recurring event is a loss of *TP53* function, which otherwise prevents the propagation of CIN cells [18].

The CIN 70 signature is a set of genes whose expression is correlated with functional segmental aneuploidy [1]. It was one of the first CIN signatures and it is enriched by genes involved in cell cycle regulation and mitosis. CIN 70 was later criticised for rather being a marker for cell proliferation than for CIN, because it reflects evolved aneuploid cancer cell populations which have adapted their genome instead of a primary response to CIN [19]. These studies highlighted that we have to distinguish between acute responses to aneuploidy and CIN [11], mechanisms for tolerating CIN [17] and the cellular programme [20, 21] and genetic alterations [22] acquired by evolved CIN cells. These cellular programmes might differ between cancer cell lines and tumours, partially as a result of treatment effects or as a result of interactions with the tumour microenvironment. Recently, it was discovered that chromosome segregation errors as well as replication stress activate the anti-viral immune *cGAS-STING* pathway, which responds to genomic double-stranded DNA in the cytosol [2, 23]. This interesting research links cancer cell intrinsic processes with cell to cell communication and immune response in the tumour microenvironment.

The phenotypic plasticity in combination with tumour heterogeneity enables CIN tumours to rapidly adapt to diverse stress conditions. It has been shown that CIN permits and accelerates the acquisition of resistance against anticancer therapies by acquiring recurrent copy number changes [24, 25]. This acquired drug resistance could potentially exacerbate the intrinsic drug resistance [26] of many CIN cells, which highlights the need to better understand genomic changes of CIN tumours in the context of anti-cancer treatment.

Computational studies of cancer genomic data have provided valuable insights into CIN [1, 19–22] and aneuploidy [27] and guided experimental and clinical testing. However, most of these studies did not differentiate between W-CIN and S-CIN. Here, we analyse cancer genomic data to better understand commonalities and differences between both types of CIN. In particular, we analyse, across multiple cancer types, the genomic landscape of S-CIN and W-CIN, their relationship to prognosis and drug sensitivity, the relationship between CIN, somatic point mutations and specific copy number variations and propose a new link between S-CIN and the *PI3K* oncogenic pathway.

## 3. Materials and Methods

### 3.1. TCGA Pan-Cancer Clinical and Molecular Data

We analysed chromosome instability of 33 primary tumour types from The Cancer Genome Atlas (TCGA): Adrenocortical carcinoma (ACC, *n* = 89); bladder urothelial carcinoma (BLCA, *n* = 399); breast invasive carcinoma (BRCA, *n* = 1039); cervical and endocervical cancers (CESC, *n* = 294); cholangiocarcinoma (CHOL, *n* = 36); colon adenocarcinoma (COAD, *n* = 420); lymphoid neoplasm diffuse large B-cell lymphoma (DLBC, *n* = 47); esophageal carcinoma (ESCA, *n* = 162); glioblastoma multiforme (GBM, *n* = 556); head and neck squamous cell carcinoma (HNSC, *n* = 510); kidney chromophobe (KICH, *n* = 65); kidney renal clear cell carcinoma (KIRC, *n* = 480); kidney renal papillary cell carcinoma (KIRP, *n* = 280); acute myeloid leukaemia (LAML, *n* = 124); brain lower grade glioma (LGG, *n* = 506); liver hepatocellular carcinoma (LIHC, *n* = 361); lung adenocarcinoma (LUAD, *n* = 490); lung squamous cell carcinoma (LUSC, *n* = 482); mesothelioma (MESO, *n* = 81); ovarian serous cystadenocarcinoma (OV, *n* = 550); pancreatic adenocarcinoma (PAAD, *n* = 165); pheochromocytoma and paraganglioma (PCPG, *n* = 160); prostate adenocarcinoma (PRAD, *n* = 471); rectum adenocarcinoma (READ, *n* = 154); sarcoma (SARC, *n* = 244); skin cutaneous melanoma (SKCM, *n* = 104); stomach adenocarcinoma (STAD, *n* = 427); testicular germ cell tumours (TGCT, *n* = 133); thyroid carcinoma (THCA, *n* = 463); thymoma (THYM, *n* = 106); uterine corpus endometrial carcinoma (UCEC, *n* = 512); uterine carcinosarcoma (UCS, *n* = 56); uveal melanoma (UVM, *n* = 80).

We also calculated karyotypic complexity scores as surrogate measures for CIN (see Section 3.4) for 391 metastatic tumour tissues, 8719 blood-derived normal tissues and 2207 solid normal tissues.

The TCGA pan-cancer molecular and clinical data were downloaded from the Pan-Cancer Atlas [28]. The file names for different data modalities are: Copy number segment data from broad.mit.edu_PANCAN_Genome_Wide_SNP_6-whitelisted.seg; ABSOLUTE [29] inferred ploidy data from TCGA mastercalls.abs t ables JSedit.fixed.txt; normalised and batch effect-corrected gene expression profile from EBPlusPlusAdjustPANCAN_IlluminaHiSeq_RNASeqV2.geneExp.tsv; clinical data from TCGA-CDR-SupplementalTableS1.xlsx; PARADIGM [30] inferred pathway activity data from merge_merged_reals.tar.gz.

### 3.2. CCLE Molecular and Sample Annotation Data

Cell line multiomics data were downloaded from the Broad-Novartis Cancer Cell Line Encyclopedia (CCLE) [31]. In particular, the copy number segment data are located in CCLE_copynumber_2013-12-03.seg.txt. Gene expression profiles and sample annotations are located in CCLE_RNAseq_genes_rpkm 201 80929.gct.gz and Cell_lines_annotations_20181226.txt. The binary alteration matrix is located in CCLE_MUT_CNA_AMP_DEL_binary_Revealer.gct. Sample ploidy data estimated using the ABSOLUTE algorithm [29] are located in CCLE_ABSOLUTE_combined_20181227.xlsx.

### 3.3. CTRP Drug Screening Data

We collected cell line pharmacological profiling data from the Cancer Therapeutics Response Portal (CTRP [32], CTRPv2.0_2015_ctd2_ExpandedDataset.zip). The drug resistance quantified by the area under the dose–response curve (AUC) was min–max normalised, i.e., the minimum value was subtracted and the resulting values were rescaled by the original range of the AUC. These min–max normalised AUC values have a range between zero and one. From this, we computed the drug sensitivity index as 1 normalised AUC with values in the range between 0 (highest resistance) and 1 (most sensitive).

### 3.4. Karyotypic Complexity Scores (CIN Scores)

We implemented three different karyotypic complexity scores [7] as surrogate measures for CIN in both TCGA bulk tumours and CCLE cell lines: The numerical complexity score (NCS), the structural complexity score (SCS) and the weighted genome instability index (WGII). For brevity, we will refer to these karyotypic complexity scores as CIN scores. Here, we detail the procedures for computing each score.

The NCS is calculated by the following steps:

Step 1: Inferring sample ploidy using the ABSOLUTE algorithm [29].

Step 2: Rounding the ploidy and segment-wise copy numbers of each sample to the nearest integer.

Step 3: Identifying whole chromosomal changes in each chromosome. For each chromosome in a sample, this chromosome is counted as a whole chromosomal change if at least 75% of the chromosome has integer copy numbers greater or less than the sample integer ploidy.

Step 4: Summing up the whole chromosome changes across all 22 autosomes yields the sample NCS.

The SCS is calculated by the following steps:

Step 1: Rounding the segment-wise copy numbers of each sample to the nearest integer.

Step 2: Computing the modal copy number for each chromosome in each sample.

Step 3: Identifying intra-chromosomal changes for each chromosome. Given a chromosome segment of a sample, this segment (with length ≥1 Mb) is counted as changed if its integer copy number is greater or less than the modal copy number of this chromosome.

Step 4: Summing up all intra-chromosomal changes across all 22 autosomes yields the sample SCS.

The WGII is calculated by the following steps:

Step 1: Inferring sample ploidy using the ABSOLUTE algorithm [29].

Step 2: Rounding the ploidy and segment-wise copy numbers of each sample to the nearest integer.

Step 3: Identifying chromosome changes for each chromosome. Given a chromosome segment of a sample, this segment is counted as changed if the integer copy number of this segment is greater or less than the sample integer ploidy.

Step 4: Calculating the percentage of the chromosome change for each chromosome.

Step 5: Calculating the mean percentage of the chromosome change of all 22 autosomes, resulting in sample WGII.

### 3.5. Association Analysis between CIN and Genome Instability

Aneuploidy scores (ASs) of samples are taken from [27], Supplementary Table S2, tumour characteristics including homologous recombination deficiency (HRD), silent mutation rate (SMR), non-silent mutation rate (NSMR), proliferation and intra-tumour heterogeneity (ITH) were collected from [33], Supplementary Table S1. Microsatellite instability (MIN) scores are collected from [34], Supplementary Table S5. The correlations of these genome instability scores and NCS or SCS were quantified by Spearman correlation coefficients.

### 3.6. Survival Analysis

We performed survival analysis using the survival R package [35]. Patients were stratified according to their median CIN score of all patients from the same cohort. A univariate Cox proportional hazards model was fitted to evaluate the association between patient survival and CIN and the log rank test was applied to calculate the *p*-value for the survival difference between high-CIN and low-CIN groups. Survival curves were visualised using ggsurvplot implemented in the survminer R package [36].

### 3.7. Treatment Response Analysis

We labelled patients with complete/partial response to chemotherapy or radiation therapy as responders and the other patients as non-responders. A Wilcoxon rank sum test was used to evaluate the differences of the NCS and SCS in the responder and non-responder groups.

### 3.8. Identification of Candidate Compounds Selectively Targeting CIN

Spearman correlation coefficients between drug sensitivity (defined in Section 3.3) and CIN were computed for 545 CTRP compounds. Compounds with multiple testing adjusted *p* ≤ 0.05 and median drug sensitivity *>*0.5 were considered as candidate compounds selectively targeting low-CIN cancer cells (compounds with negative correlation coefficients) or high-CIN cancer cells (compounds with positive correlation coefficients).

### 3.9. Association Analysis between CIN and PARADIGM Pathway Activities

We collected the sample-wise PARADIGM pathway activity matrix from the Pan-Cancer Atlas [28] with the file name merge merged reals.tar.gz. For each cancer type we computed the Spearman correlation coefficient between CIN score (NCS or SCS) and PARADIGM pathway activity and selected the top pathways corresponding to significant protein coding genes. We filtered genes/proteins whose PARADIGM pathway activities are strongly positively correlated with NCS or SCS (correlation coefficient ≥0.3) in more than seven cancer types.

### 3.10. Association Analysis between Somatic Alterations and CIN

We used the limma R package [37] for multiple linear regression analysis on CIN scores, using alteration status (mutation, copy number amplification or copy number deletion versus wild type) and cohort as predictor variables. To achieve sufficient statistical power, only alterations which occurred in more than 20 samples were included as predictors.

## 4. Results

### 4.1. Karyotypic Complexity Scores as Surrogate Measures for CIN

CIN is a dynamic feature of abnormal chromosomes, rendering its assessment in routine experimental settings difficult [38, 39]. Assessing the degree of ongoing W-CIN or S-CIN requires time-resolved data to monitor the rate of mitotic errors or the rate of segmental gains or losses, respectively. An alternative is to use single cell analysis to quantify cell to cell karyotype heterogeneity within a population of cells. The latter approach is based on the assumption that the degree of CIN is reflected by the degree of karyotype heterogeneity.

Although these and other approaches have made considerable progress in recent years (see, e.g., [40] for a recent review), the number of patient-derived tumour samples across different cancer types providing such information is not sufficient for a statistically meaningful comparison across different cancer types. Instead, we use established karyotypic complexity scores which have been evaluated as good markers for the CIN phenotype [7, 26]. Please note, however, that these scores derived from cross-sectional tumour data quantify the degree of aneuploidy or segmental aneuploidy, which is the result of both CIN and the selective pressures shaping the karyotype. As such, the karyotypic complexity scores cannot quantify ongoing CIN, but only reflect the chromosomal changes resulting from CIN and evolutionary adaptation and selection. Nevertheless, based on previous evidence [26] we assume here that these karyotypic complexity scores reflect features of the evolved CIN phenotype and refer to them as CIN scores.

As a surrogate score for the degree of W-CIN of a given tumour sample, we used the numerical complexity score (NCS) [7], which counts the number of whole chromosome gains/losses (defined as chromosomes with more than 75% of integer copy numbers higher or lower than the sample integer ploidy). The exact computation is given in Section 3.4. The degree of S-CIN was assessed by the structural complexity score (SCS), which is the number of structurally aberrant regions in the genome of a sample. A region in a chromosome is defined as structurally aberrant if it is longer than 1 Mb and its copy number deviates from the modal copy number of the chromosome (Section 3.4).

The weighted genome instability index (WGII) was previously used as a measure integrating both numerical and structural complexity (e.g., [7, 12]).

The WGII is the average percentage of changed genome relative to the sample ploidy [7], see again Section 3.4. We found that the WGII is highly correlated to the NCS (Pearson correlation coefficient: 0.99) and we also provide the pan-cancer analysis results using the WGII for comparison in PDF S1.

Please note one important difference between our work and previous analysis (e.g., [7, 12]) of karyotypic complexity scores: We used the ABSOLUTE algorithm for estimating the ploidy of the sample, whereas most previous work used the median copy number weighted by segment length across all segments [7]. The ABSOLUTE inferred ploidy has been validated using fluorescence-activated cell sorting, spectral karyotyping and DNA-mixing experiments [29].

### 4.2. Landscape of W-CIN and S-CIN across Human Cancers

In total, we calculated NCS and SCS for 21,633 samples including 10,308 primary tumours, 391 metastatic tumours and 10,934 normal tissues derived from 33 cancer types. The distribution of NCS varies drastically across cancer types (Figure 1A), but shows a characteristic bimodal pattern, see also the pan-cancer histogram on the right hand side. The colour coding of the whole genome doubling (WGD) status indicates that tumour samples with high levels of NCS are often characterised by a WGD event. Please note that this is not an artefact of the NCS, which is measured relative to the sample ploidy. This suggests that WGD is an important mechanism inducing W-CIN in many cancer types. However, the exception is kidney chromophobe (KICH), where WGD events seem to be rare, but high levels of the NCS can still be observed. In this cancer type, there is also no clear bimodal pattern, suggesting that mechanisms other than WGD drive W-CIN in KICH. Even in cancers where the bimodal pattern suggests a clear separation between numerically unstable and numerically stable tumours, it is difficult to define a universal NCS threshold distinguishing numerically stable from W-CIN tumours across cancer types. For example, in ovarian serous cystadenocarcinoma (OV), one can distinguish low- and high-NCS groups with WGD, but the overall level of the NCS is much higher than that in other cancer types. Similarly, for adrenocortical carcinoma (ACC), there are many patients with high levels of NCS even in the group of samples which did not undergo WGD. This suggests that processes other than WGD can drive a certain degree of W-CIN in these tumours.

**Figure 1.**
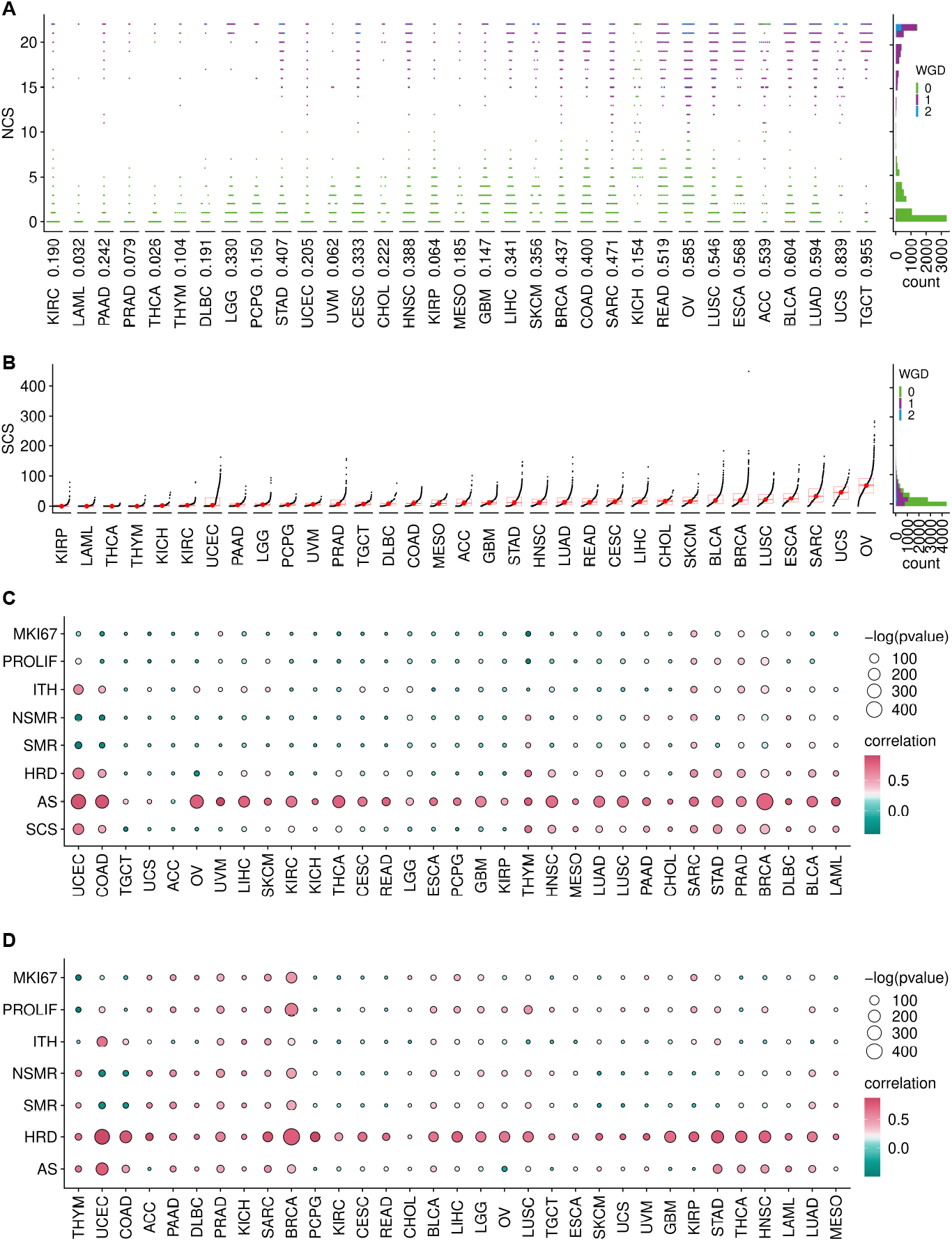
Distribution of CIN scores and their association with genetic instability. (**A**) Left: NCS for TCGA tumour samples (dots) from different cancer types, sorted according to median NCS. The colour coding indicates the WGD status and the number below each beeswarm plot is the proportion of samples which underwent WGD. Right: Pan-cancer histogram of NCS. (**B**) SCS for tumour samples from different cancer types, ordered by their median SCS. Right: Pan-cancer histogram of SCS with colours indicating WGD status. (**C**,**D**) Correlation between NCS (**C**) or SCS (**D**) with different indices for genetic instability, intra-tumour heterogeneity and proliferation: *MKI67* expression, proliferation rates (PROLIF), intra-tumour heterogeneity (ITH), non-silent mutation rate (NSMR), silent mutation rate (SMR), homologous recombination deficiency (HRD) and aneuploidy score (AS). Data for these indices were collected from [27, 33].

In contrast to the NCS distribution, the pan-cancer distribution of SCS peaks at low values and is right skewed (Figure 1B). This indicates that most tumours are structurally chromosomally stable, but some can exhibit extreme levels of S-CIN. Overall, there is no functional relationship between NCS and SCS (Figure A1).

The distribution of SCS indicates a high degree of tumour heterogeneity within the same cancer type and across cancer types. Ovarian serous cystadenocarcinoma (OV), uterine carcinosarcoma (UCS) and sarcoma (SARC) show the highest SCS (Figure 1B) and many samples within these tumours also exhibit high NCS (compare Figure 1A). Both types of CIN occur in many OV, esophageal carcinoma (ESCA) and BRCA samples, whereas thyroid carcinoma (THCA), thymoma (THYM) and acute myeloid leukaemia (LAML) samples are typically both structurally and numerically stable. Cancer types previously recognised as those dominated by the CIN phenotype [41], including stomach adenocarcinoma (STAD), colon adenocarcinoma (COAD), uterine corpus endometrial carcinoma (UCEC), OV, UCS and prostate adenocarcinoma (PRAD) have extremely heterogeneous SCS.

We also checked for associations of CIN with other types of genetic instability by correlating the NCS and SCS with different features: Aneuploidy score (AS), homologous recombination deficiency (HRD), silent mutation rate (SMR), nonsilent mutation rate (NSMR) and intra-tumour heterogeneity (ITH). The NCS is positively associated with the aneuploidy score (Figure 1C) across cancer types [42]. HRD is consistently positively associated with the SCS (Figure 1D), suggesting that impaired repair of double-strand DNA breaks might be a key driver of S-CIN.

CIN and microsatellite instability (MIN) are usually considered mutually exclusive [38]. Indeed, most MIN tumours have low NCS and SCS, but some MIN samples which underwent WGD can also exhibit signs of W-CIN and S-CIN (Figure A2A).

To check for a potential link between CIN and proliferation, we used a proliferation index [33] and the expression of the *MKI67* marker for proliferation. In many cancers, including BRCA, SARC, STAD and PRAD, increasing levels of NCS go along with increasing levels of these proliferation markers (Figure 1C). Proliferation markers are also associated with SCS in some cancers, including BRCA and LUSC. However, this is not the case for many other cancers, reflecting again the complex relationship between CIN and proliferation [43–45]. The balance between the proliferation-promoting effect of CIN as a template for Darwinian selection and the cellular burden of chromosomal aberrations accompanied by CIN might be highly cancer type dependent.

Both NCS and SCS tend to be higher in primary tumours than in normal samples (Figure A2B). Previous findings linked CIN and metastasis [23]. We find that metastatic tumours tend to have higher levels of the SCS. For the NCS, this relationship is unclear. The average NCS is higher in metastatic tumours, but there are many primary tumours with high levels of NCS. The small sample size for metastatic tumours prevents a cancer type-specific analysis of the relationship between CIN and metastatic disease.

These results highlight that W-CIN and S-CIN are two related but distinct phenotypes with different distributions across cancer types. Whole genome doubling is often accompanied by W-CIN, but this does not completely explain the elevated levels of NCS in some cancer types or individual tumours. The bimodal distribution of the NCS in most cancer types separates high-W-CIN from low-W-CIN samples, but does not provide a universal threshold valid across cancer types. However, in some cancers such as OV, even the non-WGD samples can exhibit substantial levels of W-CIN. In contrast, S-CIN is a continuous trait which is strongly associated with HRD, but not with WGD. Please note that these patterns are also observed in cell lines (Figure A2C,D).

### 4.3. Clinical Significance of CIN in Different Cancer Types

To analyse the relationship between W-CIN and prognosis, we divided the tumour samples in each individual cancer type into disjoint NCS^*high*^ and NCS^*low*^ groups using the median as a threshold. For seven of the 33 cancer types, we found that NCS^*high*^ patients had a significantly shorter overall survival than patients in the NCS^*low*^ group (Figure 2A, Table A1, log rank test, *p <* 0.05). This includes BRCA, LGG, LIHC, OV, STAD, UCEC and UVM. Disease-free survival is lower in the NCS^*high*^ group for LGG, OV, PRAD and UCEC patients (Figure A3A, Table A3, log rank test, *p <* 0.05) and progression-free survival is negatively associated with high NCS in KIRC, LGG, OV, PRAD, UCEC and UVM (Figure A4A, Table A5, log rank test, *p <* 0.05).

**Figure 2.**
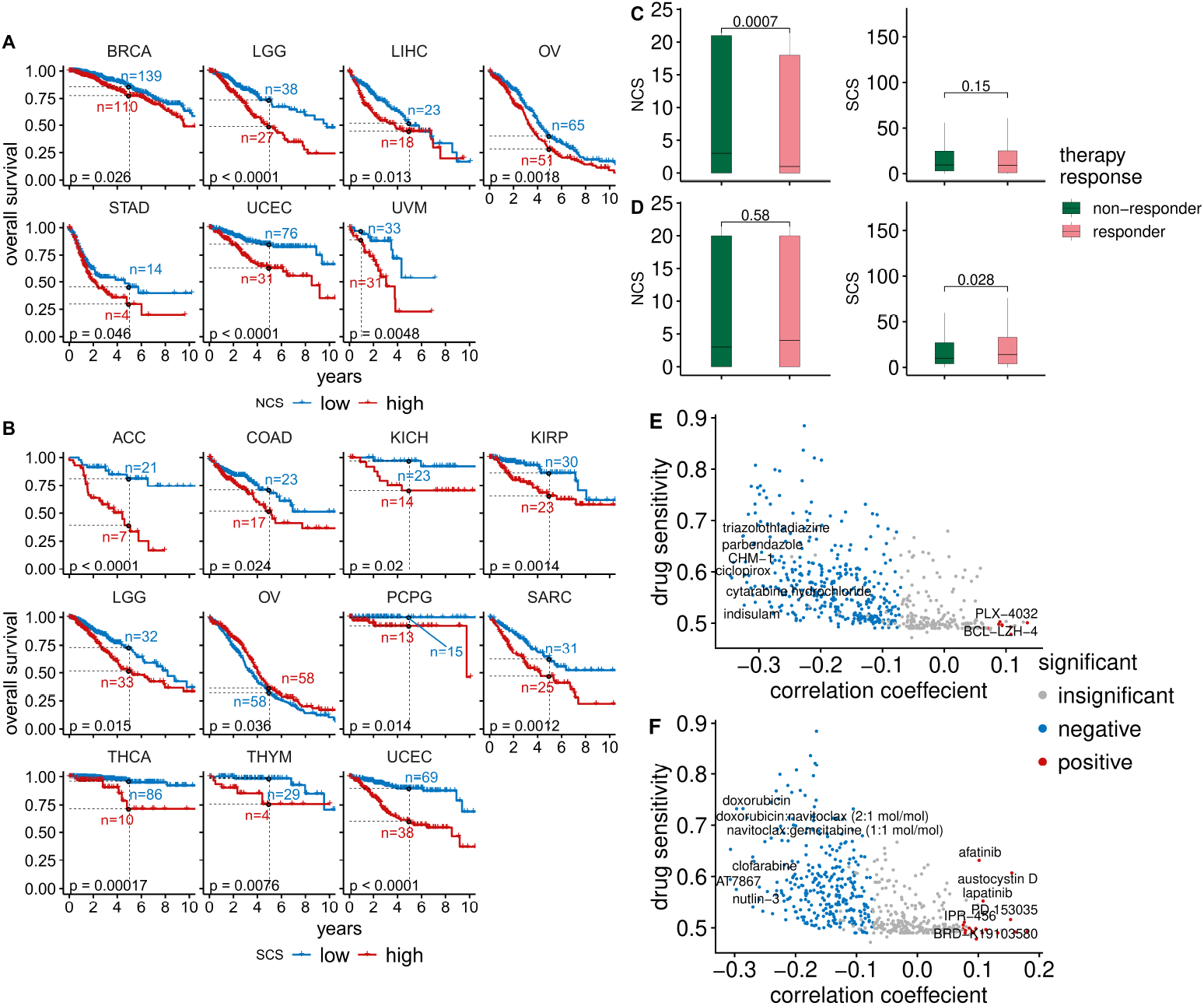
Clinical significance of CIN in different cancer types. (**A**) For seven cancer types there are significant differences in overall survival between patient samples with low NCS (blue) and high NCS (red). Dashed lines indicate the five-year overall survival probability of the two groups. (**B**) The SCS is associated with overall survival in 11 cancer types (low-SCS group in blue and high-SCS group in red). (**C**) Comparison of the NCS and SCS between radiotherapy responders and non-responders using a Wilcoxon rank sum test. (**D**) Comparison of the NCS and SCS between chemotherapy responders and non-responders using a Wilcoxon rank sum test. (**E**) The median drug sensitivity of a compound plotted against the correlation coefficient between drug sensitivity and NCS. Drugs with significant positive and negative correlations between their sensitivity and NCS are highlighted in red and blue, respectively. (**F**) The median drug sensitivity of a compound plotted against the correlation coefficient between drug sensitivity and SCS. Compounds whose sensitivity is significantly negatively or positively correlated with SCS are highlighted in blue and red, respectively.

Using an analogous separation of the tumour samples into *SCS*^*low*^ and *SCS*^*high*^ groups using the median SCS in each tumour type, we found that the overall survival of patients in 11 out of 33 cancers is negatively associated with S-CIN (Figure 2B, Table A2, log rank test, *p <* 0.05). High SCS is linked to impaired disease-free survival in adrenocortical carcinoma (ACC), KIRC, kidney renal papillary cell carcinoma (KIRP), lung squamous cell carcinoma (LUSC), PRAD, THCA and UCEC (Figure A3B, Table A4, log rank test, *p <* 0.05). For OV, patients with high SCS tend to have slightly better overall survival (Figure 2B, Table A2). However, the effect is very small and at the edge of statistical significance. In addition, the analysis of disease-free survival (Figure A3, Tables A3 and A4) and progression-free survival (Figure A4, Tables A5 and A6) does not provide any evidence for an effect of S-CIN on the prognosis of OV patients.

To further explore the clinical relevance of both types of CIN in therapy, we studied the association between CIN and response to radiotherapy or chemotherapy. Radiotherapy responders tend to have lower NCSs than radiotherapy non-responders (Wilcoxon rank test, *p* = 0.0007), whereas SCS is not significantly associated with radiotherapy response (Figure 2C). On a pan-cancer level, we did not find a significant difference between NCSs in the group of chemotherapy responders versus non-responders (Figure 2D). The median SCS of chemotherapy responders is slightly higher. One possible explanation is that high S-CIN samples tend to have defective homologous recombination repair (see Figure 1B), which renders them slightly more sensitive to chemotherapy [46, 47].

Next, we asked whether there are drugs suitable for targeting CIN [48]. To this end, we combined data from the Cancer Therapeutics Response Portal (CTRP) and the Cancer Cell Line Encyclopedia (CCLE). We normalised the area under the dose–response curve (AUC) values of 545 compounds and small molecules in all cell lines to values between zero and one and defined drug sensitivity as one minus the normalised AUC. Values of zero indicate the highest resistance level, whereas values of one indicate the highest possible sensitivity. We then computed Spearman rank correlation coefficients between the drug sensitivity of each compound with the NCS or SCS. To analyse the typical drug sensitivity as a function of CIN, we plotted the median drug sensitivity of each compound or small molecule across cell lines against their correlation coefficients with NCS (Figure 2E) or SCS (Figure 2F).

For the majority of compounds, we found negative correlations between their sensitivity and both types of CIN (Figure 2E,F), highlighting that for many compounds CIN confers an intrinsic drug resistance [26]. Only a few compounds are more potent in high-CIN cell lines than in low-CIN cell lines. However, their overall levels of sensitivity are typically low in comparison to drugs more efficient in low-CIN cell lines.

The strongest positive correlations between drug sensitivity and NCS (Figure 2E) were found for the compounds PLX-4032 and BCL-LZH-4 (median drug sensitivity *>*0.5 and FDR-adjusted *p <* 5%). PLX-4032 targets *BRAF* and has been approved by the FDA for clinical use. The *BCL2/BCL-xL/MCL1* inhibitor BCL-LZH-4 is a probe.

Drugs showing increasing sensitivity with the SCS (Figure 2F) include afatinib and lapatinib (median drug sensitivity *>*0.5 and FDR-adjusted *p <* 5%). Lapatinib targets *HER2/neu* and is used in combination treatment of *HER2*-positive breast cancer. Afatinib is used to treat non-small lung cancers with *EGFR* mutations [49]. Austocystin D is a natural cytotoxic agent and also more efficient in high-S-CIN tumours. Further details about the correlations between CIN scores and drug sensitivity can be found in the Supplementary Tables (NCS: Table S1; SCS: Table S2; WGII: Table S3).

Overall, the analysis shows that the prognostic value of CIN scores depends on cancer types and that S-CIN and W-CIN provide distinct prognostic information. The prognosis for many cancer types worsens with increased levels of CIN scores. Only for OV did we find a slightly better overall survival for patients with high SCS. It is possible that a stratification of patients according to cancer subtypes might reveal more fine-grained insights regarding the prognostic value of CIN [15, 16]. Our drug sensitivity analysis reveals that most compounds are less efficient in high-CIN tumours than in low-CIN tumours. There are a few drugs to which high-CIN cells are more sensitive than low-CIN cells. In particular, we suggest that afatinib, lapatinib and austocystin D merit further investigation for targeting S-CIN tumours. However, current drug sensitivity screens do not include many highly potent drugs specifically targeting CIN.

### 4.4. PARADIGM Pathway Activity and CIN

To identify pathways with altered activity in W-CIN or S-CIN tumours, we used the PARADIGM framework [30]. PARADIGM is a computational model which represents interactions between biological entities as a factor graph. PARADIGM integrates copy number and gene expression data and computes activities for each PARADIGM pathway feature in an individual tumour sample. These features refer to protein-coding genes, protein complexes, abstract processes and gene families. We focused on the PARADIGM features for protein-coding genes, because these are easier to interpret and can be used to generate experimentally testable predictions. We correlated the PARADIGM pathway features with the NCS or SCS and filtered features with a significant (*FDR*-adjusted *p <* 5%) Spearman correlation coefficient ≥ 0.3 in at least seven of the 32 cancer types (NCS: Figure 3A, SCS: Figure 3C).

**Figure 3.**
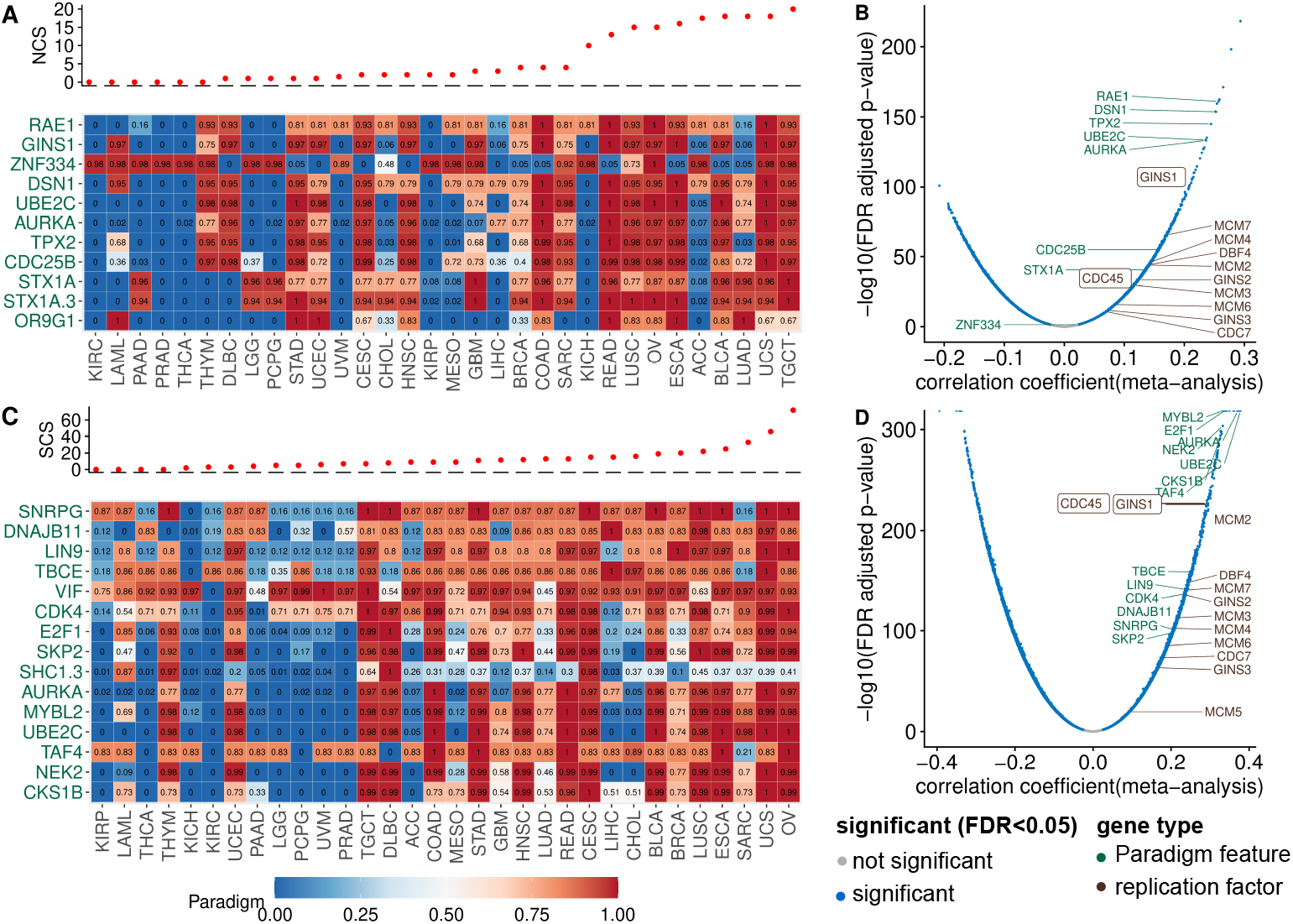
PARADIGM pathway activity and gene expression associated with CIN. (**A**) The PARADIGM pathway-level activities corresponding to protein-coding genes (rows) were correlated with the NCS. Only pathways with a significant correlation (FDR-adjusted *p <* 5%) larger than 0.3 in at least seven cancer types were included. The heatmap shows the normalised PARADIGM pathway activity (0–1 from low to high). Cancer types were ordered according to their median NCS, see top panel. (**B**) Volcano plot for the correlation between gene expression and NCS. (**C**) Analogous to (**A**), but for the SCS instead of the NCS. (**D**) Correlation of SCS and gene expression, analogous to (**B**).

PARADIGM pathway features corresponding to the mitotic genes *TPX2, RAE1, UBE2C, AURKA* (see Figure 3A) show increased activity in tumours with high NCS, consistent with the known role of chromosome segregation errors in W-CIN [1, 20]. Additionally, the PARADIGM features corresponding to the genes *CDC25B* and *DSN1* have higher activity in tumours with high NCS across many cancers. *CDC25B* regulates cell cycle progression and unregulated *CDC25B* induces replication stress, leading to CIN [50]. *DSN1* is required for kinetochore assembly.

The *STX1* (SYNTAXIN 1A) pathway shows increased activity in W-CIN tumours. This finding is surprising, because the *STX1* gene is normally expressed in brain cells and is a key molecule in synaptic exocytosis and ion channel regulation. The reason why *STX1* is upregulated in W-CIN tumours needs further investigation.

It is interesting to note the positive association of the PARADIGM feature for *GINS1* with NCS [10]. The *GINS1* protein is essential for the formation of the *Cdc45–MCM–GINS* (CMG) complex which functions to unwind DNA ahead of the replication fork [51]. As detailed in [10], overexpression of *GINS* in vitro increases replication origin firing and triggers whole chromosome missegregation and W-CIN. Indeed, when we complement our PARADIGM pathway analysis with simple gene-wise correlation of the NCS and gene expression, we find many genes involved in DNA replication and replication origin firing (see Figure 3A,B).

The analysis of the SCS-associated PARADIGM features (Figure 3C) again revealed proteins involved in kinetochore function, mitotic progression and spindle assembly and chromosome segregation (*AURKA, UBE2C, NEK2, TBCE*) or cell cycle progression (*CDK4, E2F1*).

The activity of the cyclin-dependent kinase regulatory subunit 1B (*CKS1B*) pathway is positively associated with the SCS. *CKS1B* has recently been linked to cancer drug resistance and was discussed as a new therapeutic target [52]. Our results suggest that the *CKS1B* activity is closely linked to S-CIN, which needs to be considered when studying *CKS1B* as a new target gene or as a marker of drug resistance.

To check for the robustness of these findings, we also performed a gene-wise correlation of the SCS and gene expression (Figures 3C and A5C). We also highlighted genes involved in DNA replication. Gene set enrichment analysis indicates that the top high-S-CIN-associated genes are enriched with replication origin factors (Figure A5A,B).

Please note that the analysis of genes and PARADIGM pathways negatively associated with CIN did not reveal a similarly consistent pattern across cancer types (see Figure A6).

Taken together, our analysis of PARADIGM pathway activity and gene expression in the context of CIN not only recovered known CIN genes involved in mitotic processes and spindle assembly, but highlighted, amongst others, the replication factor *GINS1* to be associated with W-CIN [10] and the *CDK* regulator and drug resistance protein *CKS1B* as strongly associated with S-CIN. In addition, we observed that the over-expression of genes involved in DNA replication is positively associated with high CIN.

### 4.5. Somatic Point Mutation Frequencies in High-CIN Tumours

To investigate the relationship between somatic point mutations and CIN, we identified genes that are more frequently or less frequently mutated in high-CIN tumours. From the 19,171 gene mutations, we included only those occurring in more than 19 samples in the wild type or mutant group across different cancer types. We fitted a linear regression model using NCS or SCS as response and somatic point mutation status (present or absent) and cancer type as predictors. The estimated regression coefficient for mutation status was used to measure its association with CIN, adjusted for tumour type.

As expected, at the pan-cancer level, *TP53* mutation shows the strongest association with CIN. Tumours harbouring a *TP53* mutation have on average more than four more whole chromosome gains or losses (ANOVA *p*-value *「* 2.2 10^*™*16^) than tumours with wild type *TP53* (Figure 4A). The mean difference in the SCS in a tumour sample with a *TP53* mutation compared to wild type samples is approximately 11 structural aberrations (Figure 4B).

**Figure 4.**
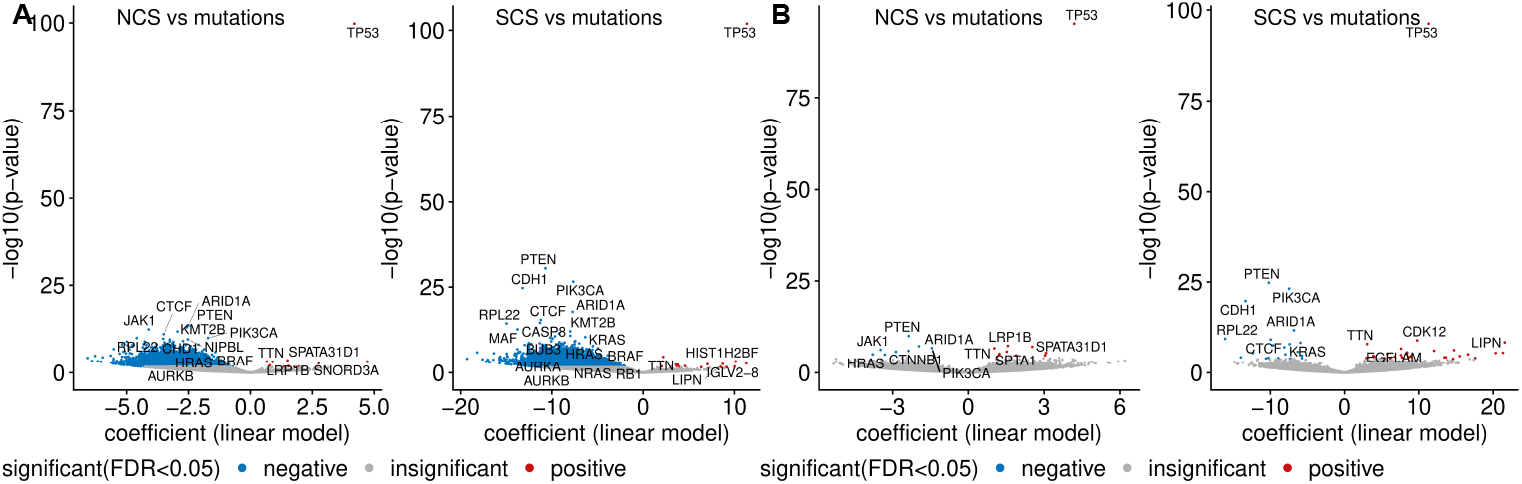
Pan-cancer somatic mutations and CIN. (**A**) The volcano plots show the association between somatic mutations and the NCS (left) or the SCS (right). The linear model coefficient indicates the mean difference of the respective CIN score when the mutation is present in a tumour sample relative to the wild type. Genes with lowest *p*-values, well-known CIN genes and cancer driver genes are highlighted. The analysis was performed on genes for which samples sizes for both wild type group and mutated group are larger than 19. Mutations significantly associated (FDR *<* 5%) with higher or lower CIN score are highlighted in blue and red, respectively. (**B**) The same as (**A**), but hypermutated MIN samples are excluded

In line with this, *TP53* mutation is positively associated with high CIN in many individual cancer types (Figure A7A). In fact, even after removing MIN samples, this correlation still holds (Figure A7B), corroborating the well-known role of *TP53* as a gatekeeper of genome stability (see e.g., [53]).

Contrary to the enrichment of *TP53* mutation in both types of CIN, we find that the presence of mutations in 5807 different genes is negatively associated with both NCS and SCS (Figure 4A). A similar negative correlation between the frequencies of recurrent copy number alterations and somatic mutations has previously been reported [54]. Later, it was realised that this negative relationship can be reversed, when the confounding effect of MIN [21, 27] is removed. When we exclude these hypermutated samples, we observe a more even distribution between genes more or less frequently mutated in high-CIN compared to low-CIN tumours (Figure 4B). This is also consistent with Figure 1C,D, where we found that neither the silent mutation rate nor the non-silent mutation rate is associated with NCS and SCS.

Intriguingly, even after excluding hypermutated samples, we find somatic point mutations of important cancer genes including *PI3KCA, PTEN* and *ARID1A* to be under-represented in high-CIN bulk tumours (Figure 4B) and high-CIN cancer cell lines (Figure A7D). *HRAS* and *JAK1* mutations are less frequent in tumours with high NCS and *KRAS* mutations are under-represented in samples with high SCS. More remarkably, when only considering validated cancer driver somatic mutations [55], the above observed relationship between *PI3KCA* mutation, *PTEN* mutation and CIN still holds (Figure A7C). The under-representation of somatic mutations in these key cancer genes in high-CIN tumours cannot be explained by differences in the overall mutation rates of these samples.

### 4.6. Copy Number Gains and Losses Associated with CIN

Given that somatic mutations of many genes are under-represented in high-CIN tumours, we next investigated copy number alterations which are specifically linked to CIN (Figure 5A). One of the strongest associations between a copy number gain and SCS was found for the *MYC* proto-oncogene. The candidate oncogene *PVT1* is also specifically gained in tumours with high SCS. *PVT1* is involved in the regulation of *MYC* [56] and carries a *TP53* -binding site. In addition, we found high NCS is associated with copy number gains for genes encoding members of the *WFDC-EPPIN* family, which have been linked to proliferation, metastasis, apoptosis and invasion in ovarian cancer (reviewed in [57]).

**Figure 5.**
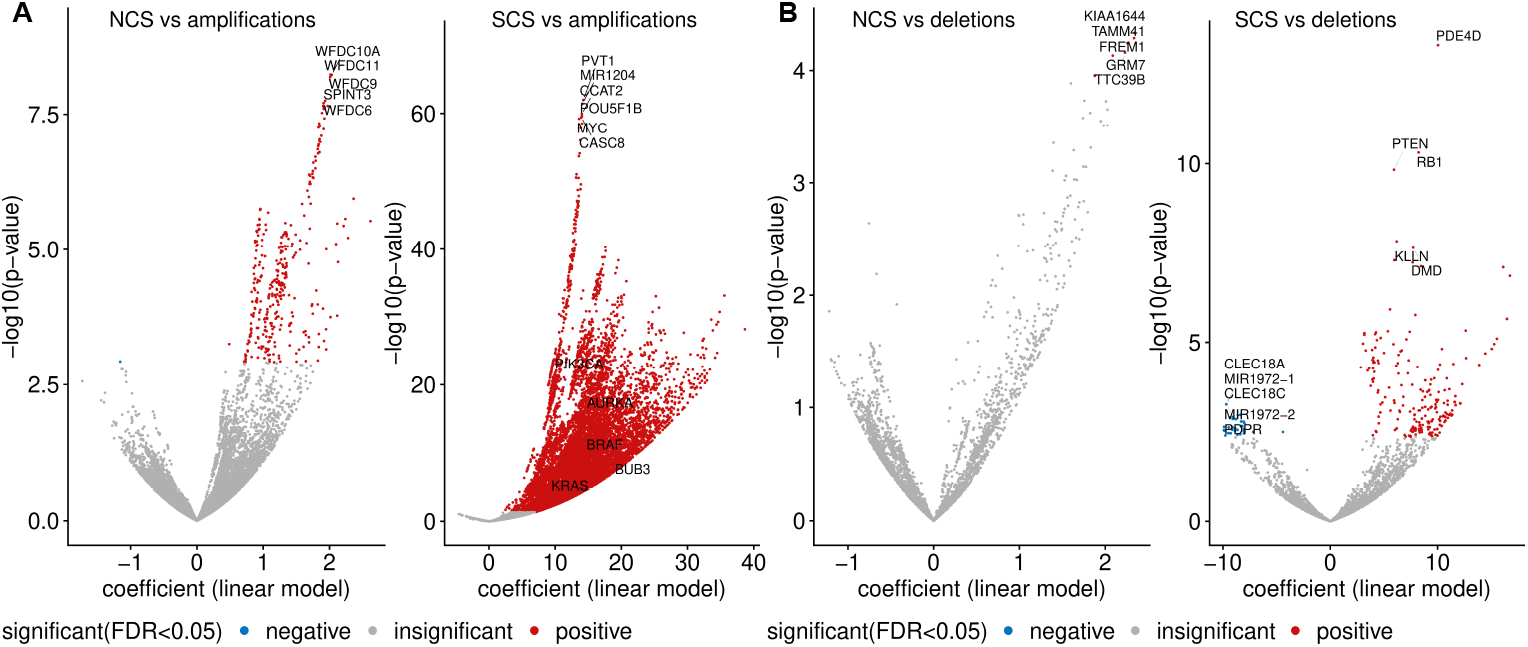
Copy number amplifications and deletions enriched in high-CIN samples. (**A**) The volcano plots show the gene-wise associations between copy number amplification status and NCS (left) and SCS (right), obtained from a regression model adjusted by cancer type. The linear model coefficient indicates the mean difference in the respective CIN score when the alteration is present in a tumour sample relative to the wild type. Genes with the lowest p-values and well-known CIN genes are highlighted. Blue and red colours encode genes with a significantly higher alteration frequency (FDR *<* 5%) in samples with low and high CIN scores, respectively. The analysis was performed on 16,922 genes with sample sizes greater than 19 for both wild type and amplified groups. (**B**) Pan-cancer copy number deletions associated with SCS are displayed in an analogous way to (**A**).

Genes specifically lost in tumour samples with high NCS include *KIAA1644, TAMM41, GRM7, TTC39B* and *FREM1* (Figure 5B). The top genes whose copy number loss is strongly associated with SCS are *PDE40, RB1* and *PTEN* (Figure 5C). The tumour suppressor *RB1* is a key regulator of the G1/S transition of the cell cycle and is required for the stabilisation of heterochromatin.

### 4.7. PI3KCA Copy Number Gains in High-S-CIN Tumours Suggest a Gene Dosage-Dependent Mechanism for PI3K Pathway Activation

In Section 4.5, we observed that somatic point mutations of *PTEN* and *PIK3CA* were scarce in high-CIN tumours. In addition, copy number amplification of *PIK3CA* and copy number loss of *PTEN* are very frequent in tumour samples with high SCS. This led us to ask whether there is a link between S-CIN and specific gene copy number alterations in these two genes to activate the *PI3K* oncogenic pathway. The *PIK3CA* gene encodes the catalytic subunit of phosphatidylinositol 3-kinase and the *PI3K* oncogenic pathway is frequently deregulated in many cancers. *PTEN* is a tumour suppressor gene and negatively regulates the growth-promoting *PI3K/AKT/mTOR* signal transduction pathway.

The oncoprint in Figure 6A displays tumour samples from all 33 TCGA cancer types in our investigation, which harbour at least one of the following genetic alterations: Somatic mutation of *PIK3CA* or *PTEN*, copy number amplification of *PIK3CA*, deletion of *PTEN*. It is apparent that there is only a small number of cancers with an amplification of *PKI3CA* or a deletion of *PTEN*, which simultaneously harbour somatic mutations in any of these genes. The copy number of both genes is also strongly associated with their gene expression. In particular, amplification and simultaneous over-expression of *PIK3CA* are associated with higher levels of SCS.

**Figure 6.**
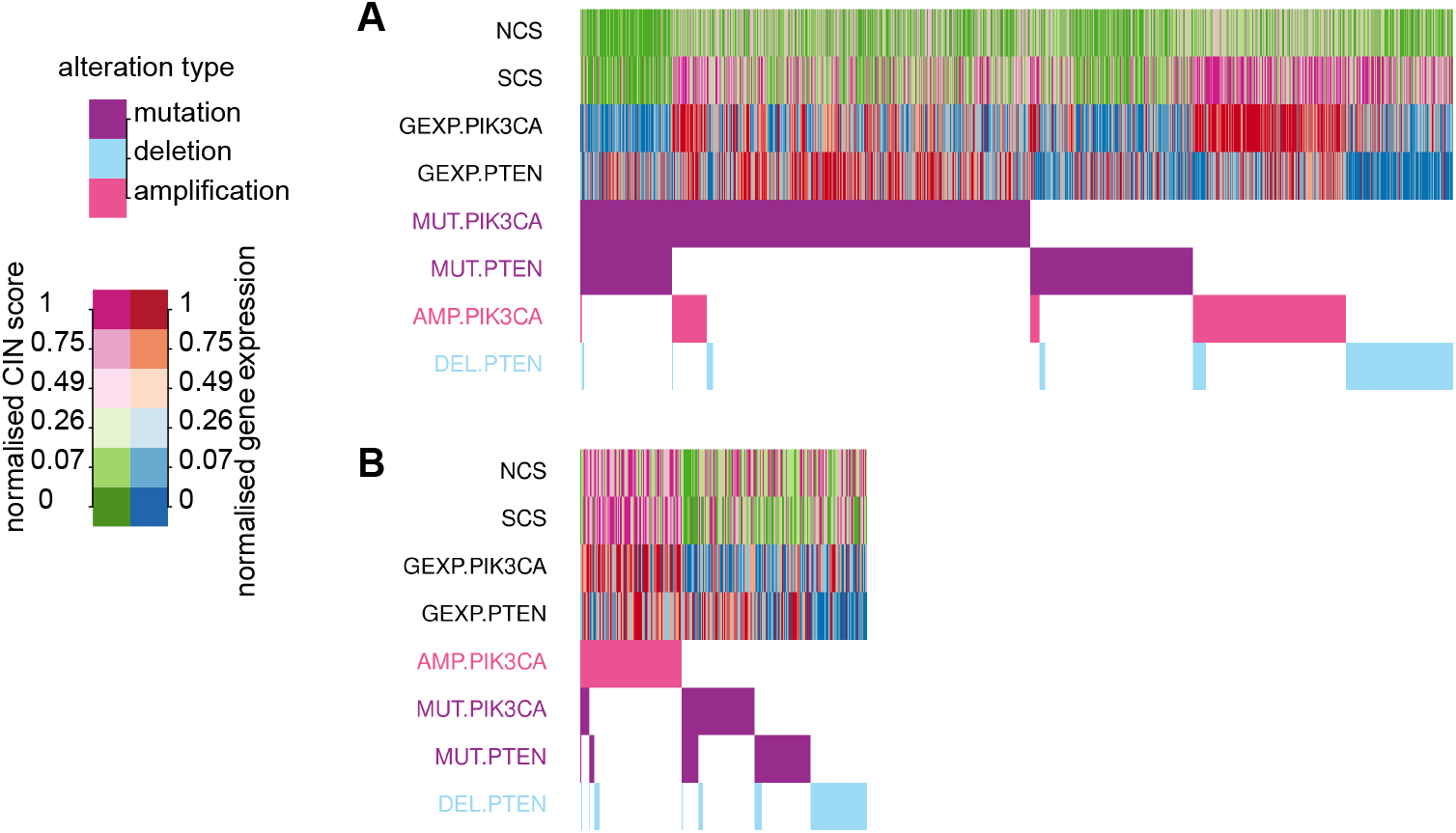
Oncoprint for *PIK3CA* and *PTEN* in relation to CIN. (**A**) The bottom panel depicts the presence or absence of somatic mutations, copy number amplifications of *PIK3CA* and deletions of *PTEN* in TCGA tumour samples (columns). Alterations are sorted by their frequency. The upper panel shows the NCS, SCS, *PI3KCA* and *PTEN* gene expression. Different levels of CIN scores and gene expression are encoded by colours. (**B**) The corresponding oncoprint for cell line data from CCLE.

To check whether this effect is preserved in pure cancer cells, we used cell line data from CCLE and found a very similar pattern. Copy number gains of *PIK3CA* are linked to high levels of its gene expression, and rarely co-occur with somatic mutations, but are associated with high SCS.

Taken together, we suggest a gene dosage effect on *PI3K* pathway activity, which is facilitated in high-S-CIN tumours. This effect is cancer cell intrinsic, because it can also be observed in cancer cell lines.

## 5. Discussion

W-CIN and S-CIN are two distinct but related phenotypes triggered by different biological mechanisms and leading to diverse consequences. A large majority of pan-cancer association studies has focused on CIN in general or exclusively on W-CIN. Here, we present an integrative statistical analysis for 33 cancer types distinguishing between W-CIN and S-CIN. We used the NCS as a proxy measure for W-CIN and the SCS to quantify the degree of S-CIN and associated these karyotypic complexity scores with various molecular and clinical features.

Our analysis reveals that the majority of tumours with high levels of NCS underwent whole genome doubling. Whole genome doubling is an early event in tumourigenesis and has been discussed as a way to rapidly accumulate numerical and structural chromosomal abnormalities and to buffer against negative effects of mutations and aneuploidy [12, 58, 59]. The results of our analysis suggest that whole genome doubling is typically accompanied by W-CIN, but not S-CIN. Instead, we find that high SCS is linked to homologous recombination deficiency, highlighting the different processes involved in these two different CIN phenotypes [6].

Although whole genome doubling is observed in many tumour samples with high levels of W-CIN, it is not sufficient to explain the elevated NCS in many tumour samples which did not undergo whole genome doubling, as most prominently observed in KICH, ACC and OV. We speculate that replication stress is an alternative mechanism for these elevated levels of W-CIN. This is based on ample evidence that replication stress can induce CIN [7, 60] and our observation that replication factors are over-expressed in tumours with high levels of W-CIN and that over-expression of the replication genes GINS1 and CDC45 can induce W-CIN [10].

We find that NCS and SCS are associated with poor prognosis in different cancer types. Only in the case of ovarian cancer did we find that high-S-CIN patients have a slightly longer overall survival, but the difference is very small and at the edge of statistical significance. In addition, we observe slightly higher NCS in patients resistant to radiotherapy. However, the relationship between CIN and prognosis is multifaceted and depends on details of the cellular physiology [3]. For instance, extreme levels of CIN in breast cancer subtypes [15, 16] were associated with better prognosis. This indicates that a subtype-specific analysis of W-CIN and S-CIN and prognosis might potentially be an interesting future project. This might also apply for the response to radiotherapy, as improved sensitivity against radiotherapy in transplanted human glioblastoma tumours has been reported [61].

From the association of NCS and SCS with in vitro drug sensitivity, it is apparent that both types of CIN are linked to intrinsic drug resistance, corroborating earlier results in colon cancer [3, 26]. However, as a new contribution we filtered small molecules and compounds for which drug sensitivity is positively associated with S-CIN or W-CIN. The drug sensitivity of a *BRAF* inhibitor, PLX-4032, is higher in cells with higher NCS. For S-CIN, this includes the approved drugs afatinib and lapatinib and the natural cytotoxic agent austocystin D. It remains to be tested whether these drugs or compounds are indeed efficient against high-CIN tumours in vivo.

In addition to well-known CIN genes including *TPX2, UBE2C* and *AURKA*, we identified a number of new candidate CIN genes and corresponding PARADIGM pathway features [30]. One interesting new finding is the chemotherapeutic drug resistance-inducing gene *CKS1B* [52], which is strongly associated with S-CIN. *CKS1B* is a cell cycle progression gene, which is discussed as a new drug target. Here, we show that *CKS1B* is over-expressed in S-CIN tumours, which might be important for the stratification of patients. We also note that the activity of the replication origin firing factor *GINS1* is linked to W-CIN, which was mechanistically verified in a recent collaboration [10]. In this context, we also found many genes involved in DNA replication to be over-expressed in tumours with high levels of W-CIN and S-CIN.

Both W-CIN and S-CIN are strongly correlated with somatic point mutation of *TP53*. We find that many copy number gains of important onogenes and loss of tumour suppressor genes [62] are strongly associated with W-CIN and S-CIN. Most strikingly, copy number gains of the oncogene *PIK3CA* and deletion of the tumour suppressor gene *PTEN* rarely occur in combination with somatic mutations in these genes. In addition, copy number gain of *PIK3CA* is linked to increased gene expression and strongly associated with S-CIN. Intriguingly, it has recently been reported that mutations in *PIK3CA* increased in vitro cellular tolerance to spontaneous genome doubling [63]. Our results, however, suggest a gene dosage effect for the activation of the *PI3K* pathway in the context of high S-CIN. This copy number-dependent activation of *PI3K* signalling was observed in both bulk tumours and cancer cell lines, indicating that it is an intrinsic property of S-CIN cells. We suggest that copy number gains of *PIK3CA* should be further investigated for both their mechanistic role in S-CIN and for their clinical implications regarding treatment strategies and patient stratification.

As a final remark, we emphasise again that our analysis is based on the karyotypic complexity scores NCS and SCS, which are averaged measures over a population of cancer cells and reflect features of the evolved W-CIN or S-CIN phenotype. As such, our analysis can stimulate new experimental work, but it cannot cover the spatio-temporal dynamics [62, 64] of tumour heterogeneity. In particular, individual chromosome changes in single cells, which still might be important drivers of cancer progression, cannot be detected by bulk data analysis [65]. We believe that the accumulation of single cell-based data from different cancer types will be essential to better understand the effect of ongoing CIN on cancer progression in the future. This will also include the testing of concepts such as karyotype coding [66], the relationship between different karyotypic states within a cellular population and the evolutionary forces shaping cancer evolution at the level of chromosome organisation.

## 6. Conclusions

In summary, our pan-cancer analysis provides insights into the distinct and common molecular, prognostic and therapeutic characteristics of W-CIN and S-CIN. Our results suggest that whole genome doubling and homologous recombination deficiency might be the most important drivers for W-CIN and S-CIN, respectively. The predictive value of W-CIN and S-CIN depends on the cancer type. We report that most of the existing compounds preferably kill low-CIN cells, but we also suggest a few compounds with increased efficiency in high-CIN cells. High activity of *CKS1B* might be a promising S-CIN target, because its expression is linked high S-CIN. We propose a new copy number-dependent mechanism for an increased activity of the oncogenic *PI3K* pathway in high-S-CIN cancer cells, which merits experimental investigation.

## Supplementary Materials

The following are available online. Table S1: The relationship between drug sensitivity and NCS, Table S2: The relationship between drug sensitivity and SCS, Table S3: The relationship between drug sensitivity and WGII, PDF S1: Pan-cancer WGII association analysis results.

## Author Contributions

X.Z.: methodology, formal analysis, investigation, software, data curation, writing, review and editing, visualization; M.K.: conceptualization, methodology, formal analysis, investigation, writing, review and editing, supervision, project administration. All authors have read and agreed to the published version of the manuscript.

## Data Availability Statement

The source code and data used to reproduce this work are available at https://github.com/mcmzxx/pancancin.

## Acknowledgments

This study used data generated by TCGA Research Network: https://www.cancer.gov/tcga and Broad-Novartis CCLE: https://sites.broadinstitute.org/ccle

## Abbreviations

The following abbreviations are used in this manuscript:

CIN: Chromosomal instability
W-CIN: Whole chromosome instability
S-CIN: Structural chromosomal instability
WGD: Whole genome doubling
WGII: Weighted genome instability index
NCS: Numerical complexity score
SCS: Structural complexity score
AS: Aneuploidy score
HRD: Homologous recombination deficiency
SMR: Silent mutation rate
NSMR: Non-silent mutation rate
ITH: Intra-tumour heterogeneity
MIN: Microsatellite instability
AUC: Area under the dose–response curve
FDA: Food and Drug Administration

## Appendix A

**Table A1.**
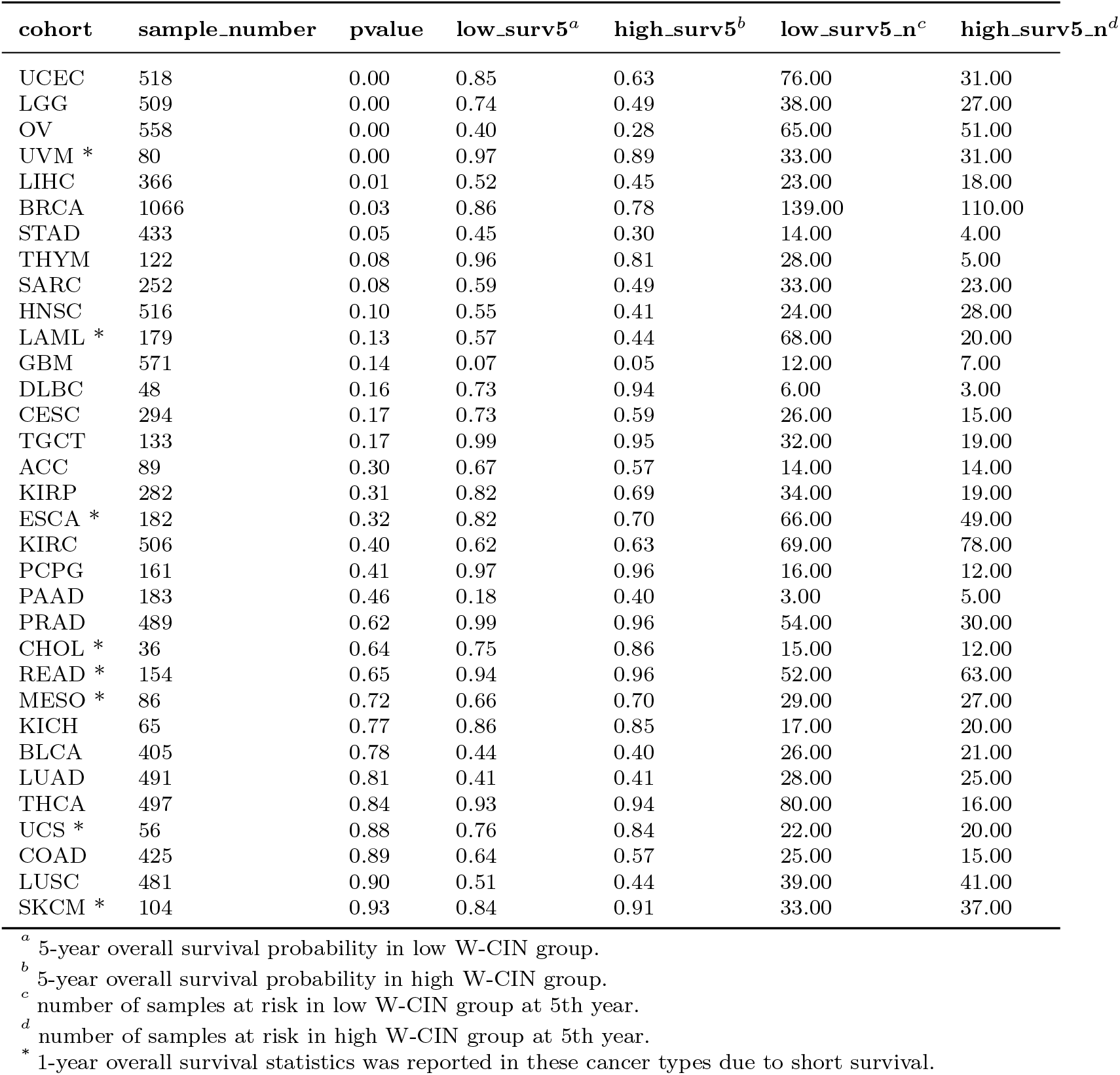
Association between W-CIN and overall survival across cancer types

| cohort | sample_number | pvalue | low_surv5<sup>a</sup> | high_surv5<sup>b</sup> | low_surv5_n<sup>c</sup> | high_surv5_n<sup>d</sup> |
| --- | --- | --- | --- | --- | --- | --- |
| UCEC | 518 | 0.00 | 0.85 | 0.63 | 76.00 | 31.00 |
| LGG | 509 | 0.00 | 0.74 | 0.49 | 38.00 | 27.00 |
| OV | 558 | 0.00 | 0.40 | 0.28 | 65.00 | 51.00 |
| UVM * | 80 | 0.00 | 0.97 | 0.89 | 33.00 | 31.00 |
| LIHC | 366 | 0.01 | 0.52 | 0.45 | 23.00 | 18.00 |
| BRCA | 1066 | 0.03 | 0.86 | 0.78 | 139.00 | 110.00 |
| STAD | 433 | 0.05 | 0.45 | 0.30 | 14.00 | 4.00 |
| THYM | 122 | 0.08 | 0.96 | 0.81 | 28.00 | 5.00 |
| SARC | 252 | 0.08 | 0.59 | 0.49 | 33.00 | 23.00 |
| HNSC | 516 | 0.10 | 0.55 | 0.41 | 24.00 | 28.00 |
| LAML * | 179 | 0.13 | 0.57 | 0.44 | 68.00 | 20.00 |
| GBM | 571 | 0.14 | 0.07 | 0.05 | 12.00 | 7.00 |
| DLBC | 48 | 0.16 | 0.73 | 0.94 | 6.00 | 3.00 |
| CESC | 294 | 0.17 | 0.73 | 0.59 | 26.00 | 15.00 |
| TGCT | 133 | 0.17 | 0.99 | 0.95 | 32.00 | 19.00 |
| ACC | 89 | 0.30 | 0.67 | 0.57 | 14.00 | 14.00 |
| KIRP | 282 | 0.31 | 0.82 | 0.69 | 34.00 | 19.00 |
| ESCA * | 182 | 0.32 | 0.82 | 0.70 | 66.00 | 49.00 |
| KIRC | 506 | 0.40 | 0.62 | 0.63 | 69.00 | 78.00 |
| PCPG | 161 | 0.41 | 0.97 | 0.96 | 16.00 | 12.00 |
| PAAD | 183 | 0.46 | 0.18 | 0.40 | 3.00 | 5.00 |
| PRAD | 489 | 0.62 | 0.99 | 0.96 | 54.00 | 30.00 |
| CHOL * | 36 | 0.64 | 0.75 | 0.86 | 15.00 | 12.00 |
| READ * | 154 | 0.65 | 0.94 | 0.96 | 52.00 | 63.00 |
| MESO * | 86 | 0.72 | 0.66 | 0.70 | 29.00 | 27.00 |
| KICH | 65 | 0.77 | 0.86 | 0.85 | 17.00 | 20.00 |
| BLCA | 405 | 0.78 | 0.44 | 0.40 | 26.00 | 21.00 |
| LUAD | 491 | 0.81 | 0.41 | 0.41 | 28.00 | 25.00 |
| THCA | 497 | 0.84 | 0.93 | 0.94 | 80.00 | 16.00 |
| UCS * | 56 | 0.88 | 0.76 | 0.84 | 22.00 | 20.00 |
| COAD | 425 | 0.89 | 0.64 | 0.57 | 25.00 | 15.00 |
| LUSC | 481 | 0.90 | 0.51 | 0.44 | 39.00 | 41.00 |
| SKCM * | 104 | 0.93 | 0.84 | 0.91 | 33.00 | 37.00 |

<sup>a</sup> 5-year overall survival probability in low W-CIN group.<sup>b</sup> 5-year overall survival probability in high W-CIN group.<sup>c</sup> number of samples at risk in low W-CIN group at 5th year.<sup>d</sup> number of samples at risk in high W-CIN group at 5th year.
\* 1-year overall survival statistics was reported in these cancer types due to short survival.

**Table A2.** Association between S-CIN and overall survival across cancer types

| cohort | sample_number | pvalue | low_surv5<sup>a</sup> | high_surv5<sup>b</sup> | low_surv5_n<sup>c</sup> | high_surv5_n<sup>d</sup> |
| --- | --- | --- | --- | --- | --- | --- |
| UCEC | 518 | 0.00 | 0.90 | 0.60 | 69.00 | 38.00 |
| ACC | 89 | 0.00 | 0.81 | 0.39 | 21.00 | 7.00 |
| THCA | 497 | 0.00 | 0.96 | 0.72 | 86.00 | 10.00 |
| SARC | 252 | 0.00 | 0.62 | 0.47 | 31.00 | 25.00 |
| KIRP | 282 | 0.00 | 0.86 | 0.65 | 30.00 | 23.00 |
| THYM | 122 | 0.01 | 0.98 | 0.76 | 29.00 | 4.00 |
| PCPG | 161 | 0.01 | 1.00 | 0.92 | 15.00 | 13.00 |
| LGG | 509 | 0.01 | 0.73 | 0.52 | 32.00 | 33.00 |
| KICH | 65 | 0.02 | 0.97 | 0.70 | 23.00 | 14.00 |
| COAD | 425 | 0.02 | 0.71 | 0.52 | 23.00 | 17.00 |
| OV | 558 | 0.04 | 0.32 | 0.37 | 58.00 | 58.00 |
| ESCA* | 182 | 0.06 | 0.81 | 0.72 | 60.00 | 55.00 |
| LUAD | 491 | 0.08 | 0.48 | 0.35 | 28.00 | 25.00 |
| TGCT | 133 | 0.09 | 0.95 | 1.00 | 25.00 | 26.00 |
| UCS* | 56 | 0.09 | 0.75 | 0.85 | 20.00 | 22.00 |
| READ* | 154 | 0.10 | 0.96 | 0.93 | 64.00 | 51.00 |
| LAML* | 179 | 0.10 | 0.58 | 0.45 | 66.00 | 22.00 |
| PAAD | 183 | 0.11 | 0.25 | 0.26 | 5.00 | 3.00 |
| KIRC | 506 | 0.17 | 0.65 | 0.59 | 83.00 | 64.00 |
| BRCA | 1066 | 0.22 | 0.83 | 0.81 | 129.00 | 120.00 |
| GBM | 571 | 0.23 | 0.05 | 0.08 | 7.00 | 12.00 |
| CHOL* | 36 | 0.39 | 0.76 | 0.83 | 13.00 | 14.00 |
| PRAD | 489 | 0.45 | 0.99 | 0.97 | 40.00 | 44.00 |
| BLCA | 405 | 0.46 | 0.40 | 0.44 | 21.00 | 26.00 |
| UVM* | 80 | 0.50 | 0.93 | 0.93 | 38.00 | 26.00 |
| HNSC | 516 | 0.56 | 0.51 | 0.44 | 27.00 | 25.00 |
| LUSC | 481 | 0.57 | 0.46 | 0.49 | 36.00 | 44.00 |
| SKCM* | 104 | 0.63 | 0.84 | 0.93 | 36.00 | 34.00 |
| MESO* | 86 | 0.64 | 0.62 | 0.75 | 28.00 | 28.00 |
| LIHC | 366 | 0.73 | 0.48 | 0.50 | 23.00 | 18.00 |
| CESC | 294 | 0.77 | 0.68 | 0.66 | 20.00 | 21.00 |
| STAD | 433 | 0.90 | 0.35 | 0.43 | 11.00 | 7.00 |
| DLBC* | 48 | 0.93 | 0.96 | 0.89 | 23.00 | 15.00 |

<sup>a</sup> 5-year overall survival probability in low S-CIN group.<sup>b</sup> 5-year overall survival probability in high S-CIN group.<sup>c</sup> number of samples at risk in low S-CIN group at 5th year.<sup>d</sup> number of samples at risk in high S-CIN group at 5th year.
\* 1-year overall survival statistics was reported in these cancer types due to short survival.

**Table A3.**
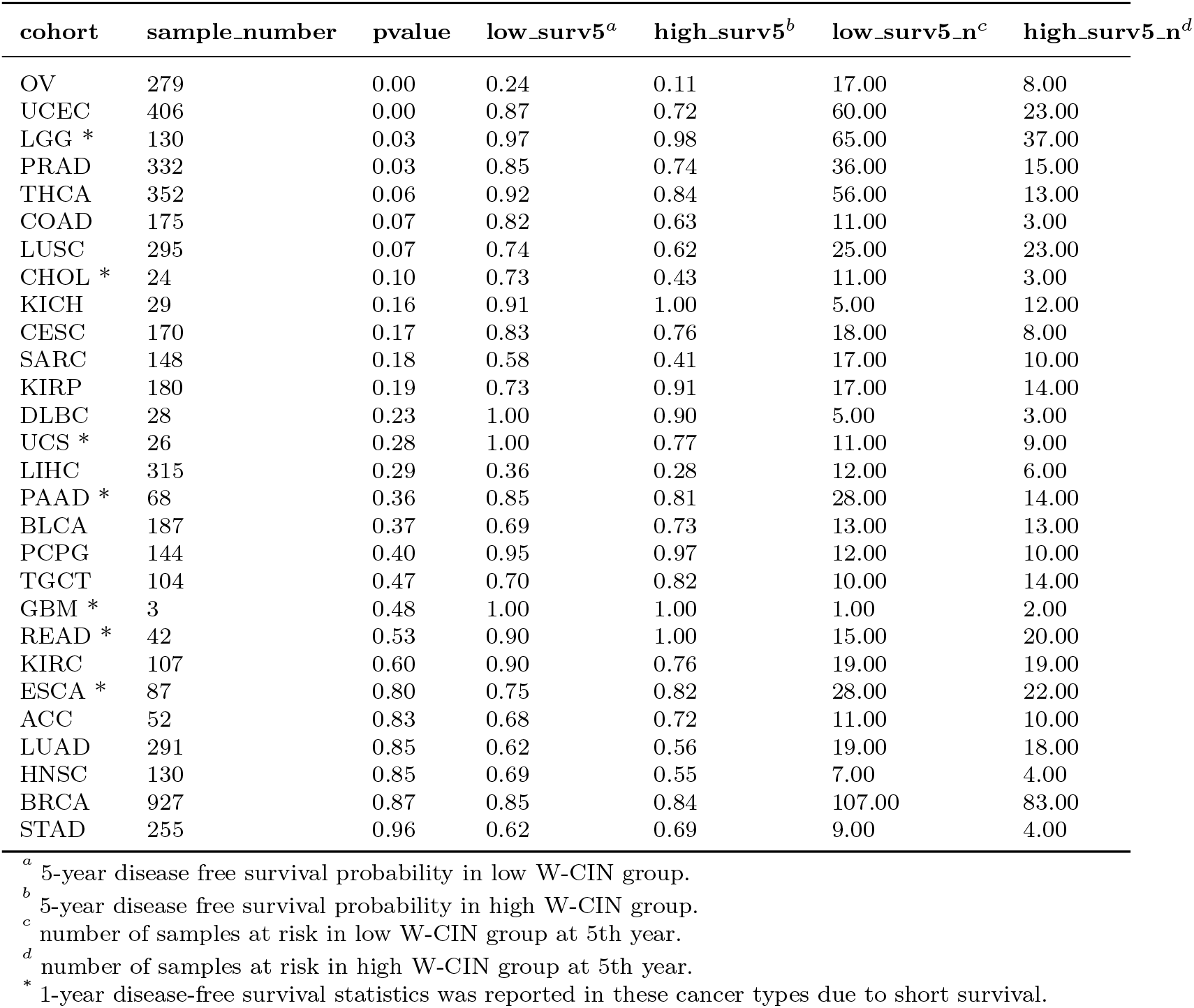
Association between W-CIN and disease free survival across cancer types

**Table A4.** Association between S-CIN and disease free survival across cancer types

| cohort | sample_number | pvalue | low_surv5<sup>a</sup> | high_surv5<sup>b</sup> | low_surv5_n<sup>c</sup> | high_surv5_n<sup>d</sup> |
| --- | --- | --- | --- | --- | --- | --- |
| UCEC | 406 | 0.00 | 0.92 | 0.65 | 61.00 | 22.00 |
| ACC | 52 | 0.00 | 0.81 | 0.48 | 16.00 | 5.00 |
| PRAD | 332 | 0.00 | 0.88 | 0.75 | 27.00 | 24.00 |
| KIRP | 180 | 0.02 | 0.88 | 0.72 | 18.00 | 13.00 |
| LUSC | 295 | 0.03 | 0.71 | 0.64 | 25.00 | 23.00 |
| THCA | 352 | 0.03 | 0.92 | 0.81 | 64.00 | 5.00 |
| KIRC | 107 | 0.04 | 0.94 | 0.70 | 24.00 | 14.00 |
| READ* | 42 | 0.09 | 1.00 | 0.89 | 20.00 | 15.00 |
| BRCA | 927 | 0.10 | 0.88 | 0.80 | 100.00 | 90.00 |
| PAAD* | 68 | 0.14 | 0.91 | 0.74 | 23.00 | 19.00 |
| LIHC | 315 | 0.15 | 0.33 | 0.31 | 12.00 | 6.00 |
| GBM* | 3 | 0.16 | 1.00 | 1.00 | 1.00 | 2.00 |
| KICH | 29 | 0.18 | 0.94 | 1.00 | 11.00 | 6.00 |
| COAD | 175 | 0.27 | 0.79 | 0.71 | 9.00 | 5.00 |
| CESC | 170 | 0.30 | 0.86 | 0.75 | 15.00 | 11.00 |
| HNSC | 130 | 0.31 | 0.66 | 0.59 | 8.00 | 3.00 |
| LUAD | 291 | 0.37 | 0.63 | 0.54 | 19.00 | 18.00 |
| DLBC* | 28 | 0.45 | 1.00 | 1.00 | 14.00 | 11.00 |
| MESO* | 15 | 0.46 | 0.69 | 1.00 | 6.00 | 2.00 |
| ESCA* | 87 | 0.49 | 0.80 | 0.76 | 26.00 | 24.00 |
| PCPG | 144 | 0.49 | 0.93 | 1.00 | 12.00 | 10.00 |
| LGG | 130 | 0.61 | 0.76 | 0.65 | 3.00 | 3.00 |
| OV | 279 | 0.64 | 0.22 | 0.14 | 17.00 | 8.00 |
| CHOL* | 24 | 0.65 | 0.67 | 0.57 | 10.00 | 4.00 |
| BLCA | 187 | 0.69 | 0.73 | 0.69 | 14.00 | 12.00 |
| TGCT | 104 | 0.70 | 0.75 | 0.75 | 14.00 | 10.00 |
| STAD | 255 | 0.76 | 0.67 | 0.64 | 8.00 | 5.00 |
| UCS* | 26 | 0.76 | 0.92 | 0.83 | 10.00 | 10.00 |
| SARC | 148 | 0.79 | 0.53 | 0.48 | 14.00 | 13.00 |

<sup>a</sup> 5-year disease free survival probability in low S-CIN group.<sup>b</sup> 5-year disease free survival probability in high S-CIN group.<sup>c</sup> number of samples at risk in low S-CIN group at 5th year.<sup>d</sup> number of samples at risk in high S-CIN group at 5th year.
\* 1-year disease-free survival statistics was reported in these cancer types due to short survival.

**Table A5.** Association between W-CIN and progression free survival across cancer types

| cohort | sample_number | pvalue | low_surv5<sup>a</sup> | high_surv5<sup>b</sup> | low_surv5_n<sup>c</sup> | high_surv5_n<sup>d</sup> |
| --- | --- | --- | --- | --- | --- | --- |
| UCEC | 518 | 0.00 | 0.80 | 0.56 | 67.00 | 27.00 |
| LGG | 509 | 0.00 | 0.53 | 0.29 | 26.00 | 10.00 |
| PRAD | 489 | 0.00 | 0.75 | 0.60 | 43.00 | 20.00 |
| UVM * | 79 | 0.00 | 0.94 | 0.69 | 31.00 | 24.00 |
| OV | 558 | 0.00 | 0.17 | 0.09 | 24.00 | 11.00 |
| KIRC | 504 | 0.04 | 0.69 | 0.58 | 60.00 | 58.00 |
| ESCA * | 182 | 0.07 | 0.66 | 0.57 | 48.00 | 36.00 |
| THYM | 122 | 0.08 | 0.83 | 0.63 | 24.00 | 5.00 |
| CESC | 294 | 0.11 | 0.69 | 0.63 | 23.00 | 13.00 |
| ACC | 89 | 0.13 | 0.49 | 0.40 | 11.00 | 10.00 |
| SKCM * | 104 | 0.15 | 0.74 | 0.68 | 27.00 | 25.00 |
| DLBC | 48 | 0.18 | 0.67 | 0.93 | 6.00 | 3.00 |
| SARC | 252 | 0.20 | 0.45 | 0.32 | 22.00 | 13.00 |
| LIHC | 366 | 0.23 | 0.29 | 0.25 | 11.00 | 7.00 |
| GBM * | 571 | 0.27 | 0.30 | 0.30 | 85.00 | 58.00 |
| LUSC | 482 | 0.31 | 0.56 | 0.53 | 32.00 | 33.00 |
| HNSC | 516 | 0.32 | 0.51 | 0.47 | 20.00 | 24.00 |
| CHOL * | 36 | 0.33 | 0.53 | 0.41 | 10.00 | 5.00 |
| THCA | 497 | 0.33 | 0.84 | 0.83 | 68.00 | 14.00 |
| READ * | 154 | 0.34 | 0.90 | 0.88 | 48.00 | 56.00 |
| PCPG | 161 | 0.40 | 0.80 | 0.88 | 11.00 | 11.00 |
| MESO * | 84 | 0.41 | 0.57 | 0.54 | 22.00 | 19.00 |
| BLCA | 406 | 0.42 | 0.39 | 0.42 | 21.00 | 15.00 |
| COAD | 425 | 0.45 | 0.62 | 0.56 | 21.00 | 9.00 |
| TGCT | 133 | 0.53 | 0.71 | 0.80 | 20.00 | 17.00 |
| STAD | 435 | 0.68 | 0.43 | 0.47 | 14.00 | 4.00 |
| KICH | 65 | 0.75 | 0.87 | 0.87 | 17.00 | 20.00 |
| BRCA | 1066 | 0.76 | 0.79 | 0.78 | 122.00 | 99.00 |
| UCS * | 56 | 0.79 | 0.48 | 0.60 | 14.00 | 14.00 |
| KIRP | 281 | 0.96 | 0.71 | 0.80 | 28.00 | 17.00 |
| LUAD | 491 | 0.99 | 0.38 | 0.40 | 20.00 | 18.00 |
| PAAD * | 183 | 1.00 | 0.64 | 0.61 | 60.00 | 32.00 |

<sup>a</sup> 5-year progression free survival probability in low W-CIN group.<sup>b</sup> 5-year progression free survival probability in high W-CIN group.<sup>c</sup> number of samples at risk in low W-CIN group at 5th year.<sup>d</sup> number of samples at risk in high W-CIN group at 5th year.
\* 1-year progression-free survival statistics was reported in these cancer types due to short survival.

**Table A6.** Association between S-CIN and progression free survival across cancer types

| cohort | sample_number | pvalue | low_surv5<sup>a</sup> | high_surv5<sup>b</sup> | low_surv5_n<sup>c</sup> | high_surv5_n<sup>d</sup> |
| --- | --- | --- | --- | --- | --- | --- |
| UCEC | 518 | 0.00 | 0.86 | 0.50 | 65.00 | 29.00 |
| ACC | 89 | 0.00 | 0.66 | 0.22 | 16.00 | 5.00 |
| KIRP | 281 | 0.00 | 0.86 | 0.61 | 27.00 | 18.00 |
| THCA | 497 | 0.00 | 0.87 | 0.64 | 75.00 | 7.00 |
| PRAD | 489 | 0.00 | 0.76 | 0.65 | 32.00 | 31.00 |
| LGG | 509 | 0.00 | 0.50 | 0.33 | 18.00 | 18.00 |
| CHOL* | 36 | 0.02 | 0.69 | 0.27 | 11.00 | 4.00 |
| THYM | 122 | 0.03 | 0.85 | 0.60 | 26.00 | 3.00 |
| GBM* | 571 | 0.06 | 0.29 | 0.32 | 73.00 | 70.00 |
| COAD | 425 | 0.07 | 0.65 | 0.53 | 19.00 | 11.00 |
| SARC | 252 | 0.08 | 0.44 | 0.34 | 19.00 | 16.00 |
| PAAD* | 183 | 0.09 | 0.67 | 0.59 | 46.00 | 46.00 |
| KICH | 65 | 0.09 | 0.97 | 0.72 | 23.00 | 14.00 |
| KIRC | 504 | 0.11 | 0.66 | 0.60 | 67.00 | 51.00 |
| ESCA* | 182 | 0.12 | 0.61 | 0.63 | 41.00 | 43.00 |
| LIHC | 366 | 0.12 | 0.26 | 0.27 | 11.00 | 7.00 |
| BRCA | 1066 | 0.12 | 0.81 | 0.75 | 118.00 | 103.00 |
| READ* | 154 | 0.12 | 0.93 | 0.84 | 59.00 | 45.00 |
| UVM* | 79 | 0.21 | 0.82 | 0.80 | 32.00 | 23.00 |
| OV | 558 | 0.25 | 0.14 | 0.13 | 22.00 | 13.00 |
| BLCA | 406 | 0.31 | 0.44 | 0.39 | 16.00 | 20.00 |
| SKCM* | 104 | 0.31 | 0.74 | 0.67 | 28.00 | 24.00 |
| LUAD | 491 | 0.35 | 0.40 | 0.38 | 19.00 | 19.00 |
| UCS* | 56 | 0.41 | 0.54 | 0.54 | 14.00 | 14.00 |
| CESC | 294 | 0.41 | 0.70 | 0.63 | 19.00 | 17.00 |
| STAD | 435 | 0.41 | 0.45 | 0.44 | 11.00 | 7.00 |
| PCPG | 161 | 0.45 | 0.84 | 0.84 | 13.00 | 9.00 |
| DLBC* | 48 | 0.51 | 0.83 | 0.84 | 19.00 | 15.00 |
| MESO* | 84 | 0.73 | 0.63 | 0.47 | 24.00 | 17.00 |
| TGCT | 133 | 0.77 | 0.72 | 0.77 | 19.00 | 18.00 |
| LUSC | 482 | 0.79 | 0.51 | 0.57 | 29.00 | 36.00 |
| HNSC | 516 | 0.93 | 0.49 | 0.49 | 23.00 | 21.00 |

<sup>a</sup> 5-year progression free survival probability in low S-CIN group.<sup>b</sup> 5-year progression free survival probability in high S-CIN group.<sup>c</sup> number of samples at risk in low S-CIN group at 5th year.<sup>d</sup> number of samples at risk in high S-CIN group at 5th year.
\* 1-year progression-free survival statistics was reported in these cancer types due to short survival.

**Figure A1.**
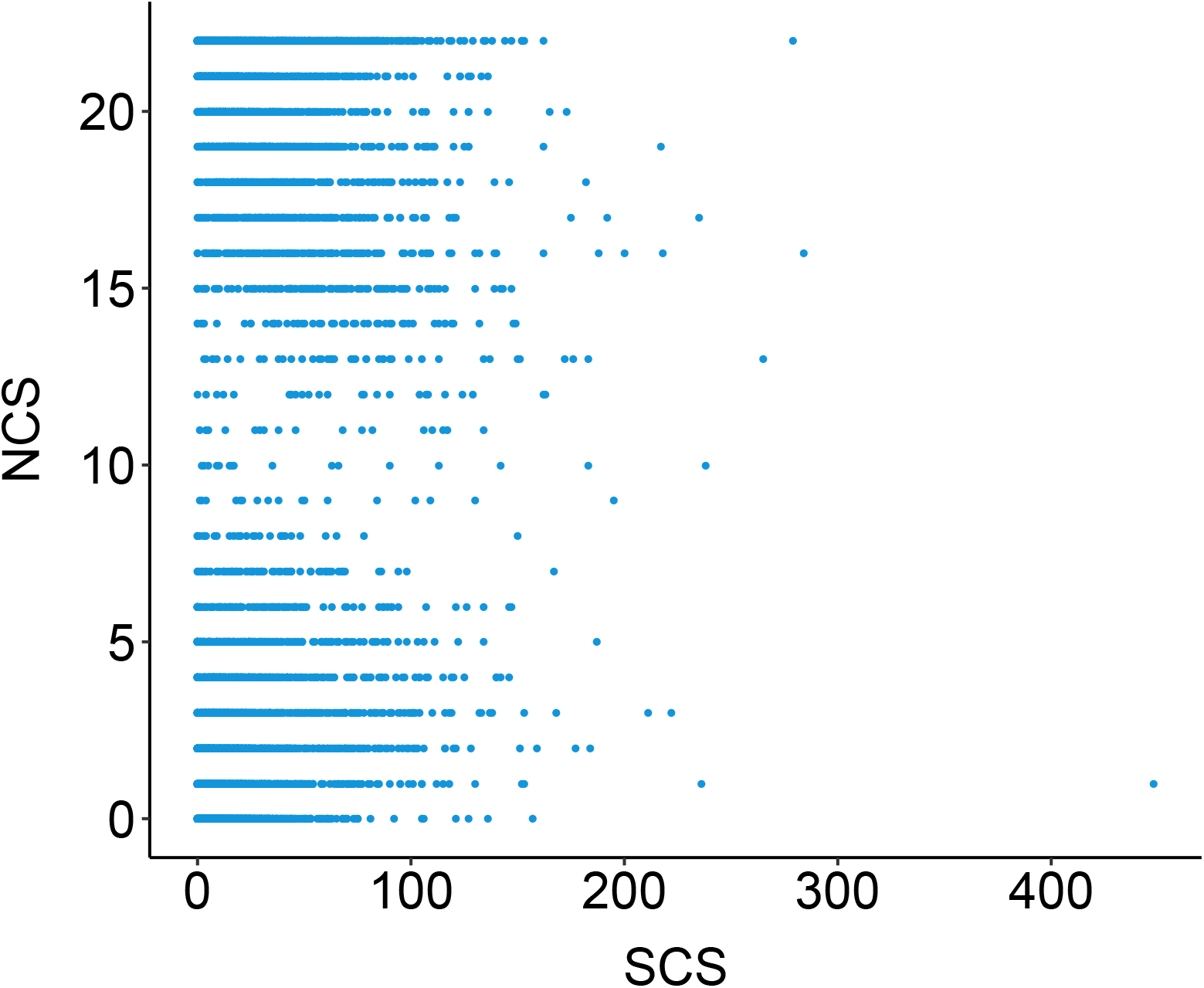
The NCS versus the SCS in TCGA tumours from 33 different cancer types.

**Figure A2.**
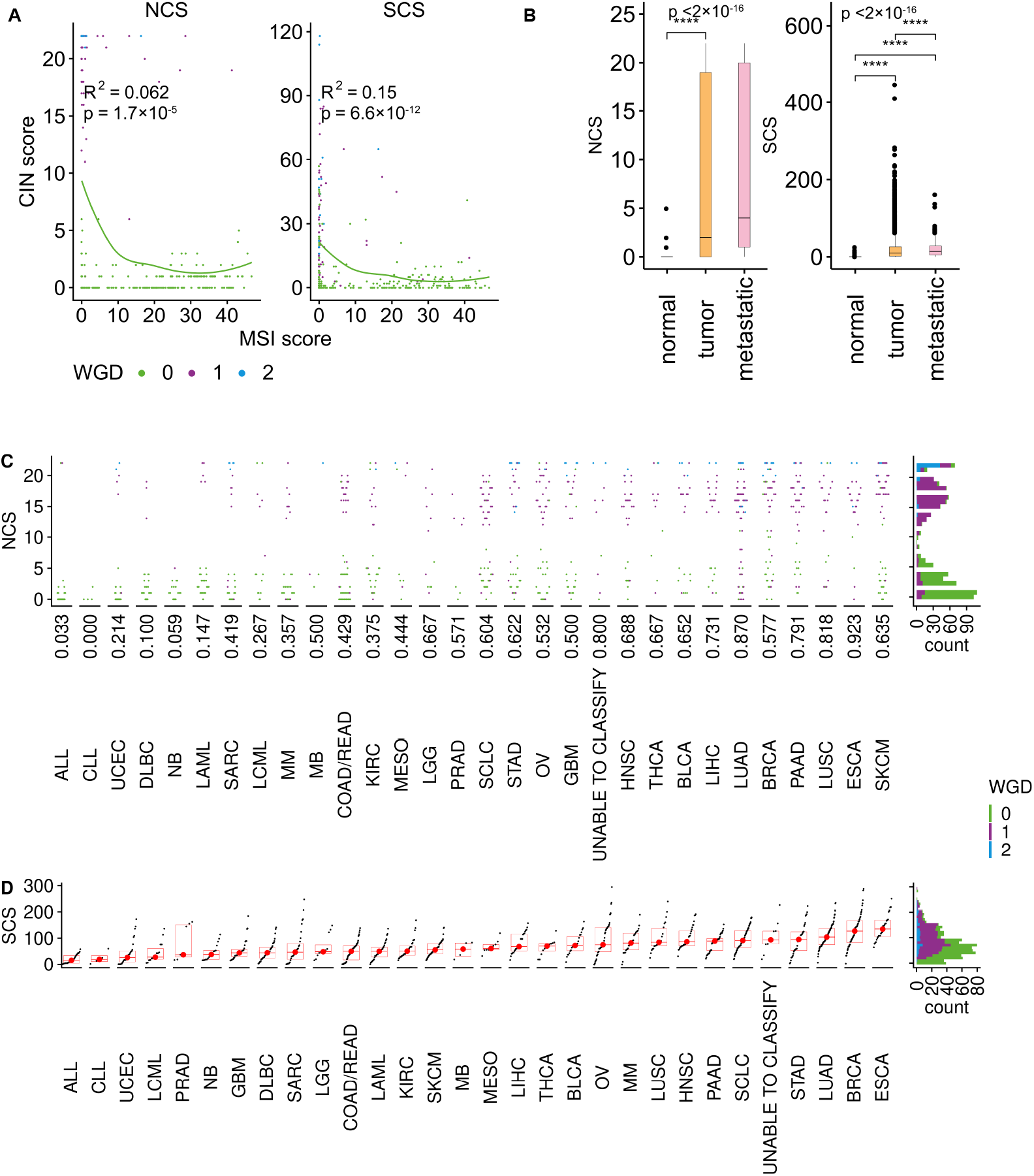
(**A**) The relationship between microsatellite instability (MIN) scores and the NCS and the SCS. (**B**) Comparison of the NCS and the SCS between normal samples with primary and metastatic tumour samples. (**C**) Cancer type-wise NCS distribution in CCLE cell lines, cancer types are ordered by the median NCS; whole genome doubling (WGD) status is encoded by colours. The number reported on x axis is the proportion of samples that underwent WGD. (**D**) SCS distribution in CCLE cell lines, cancer types are ordered by their median SCS.

**Figure A3.**
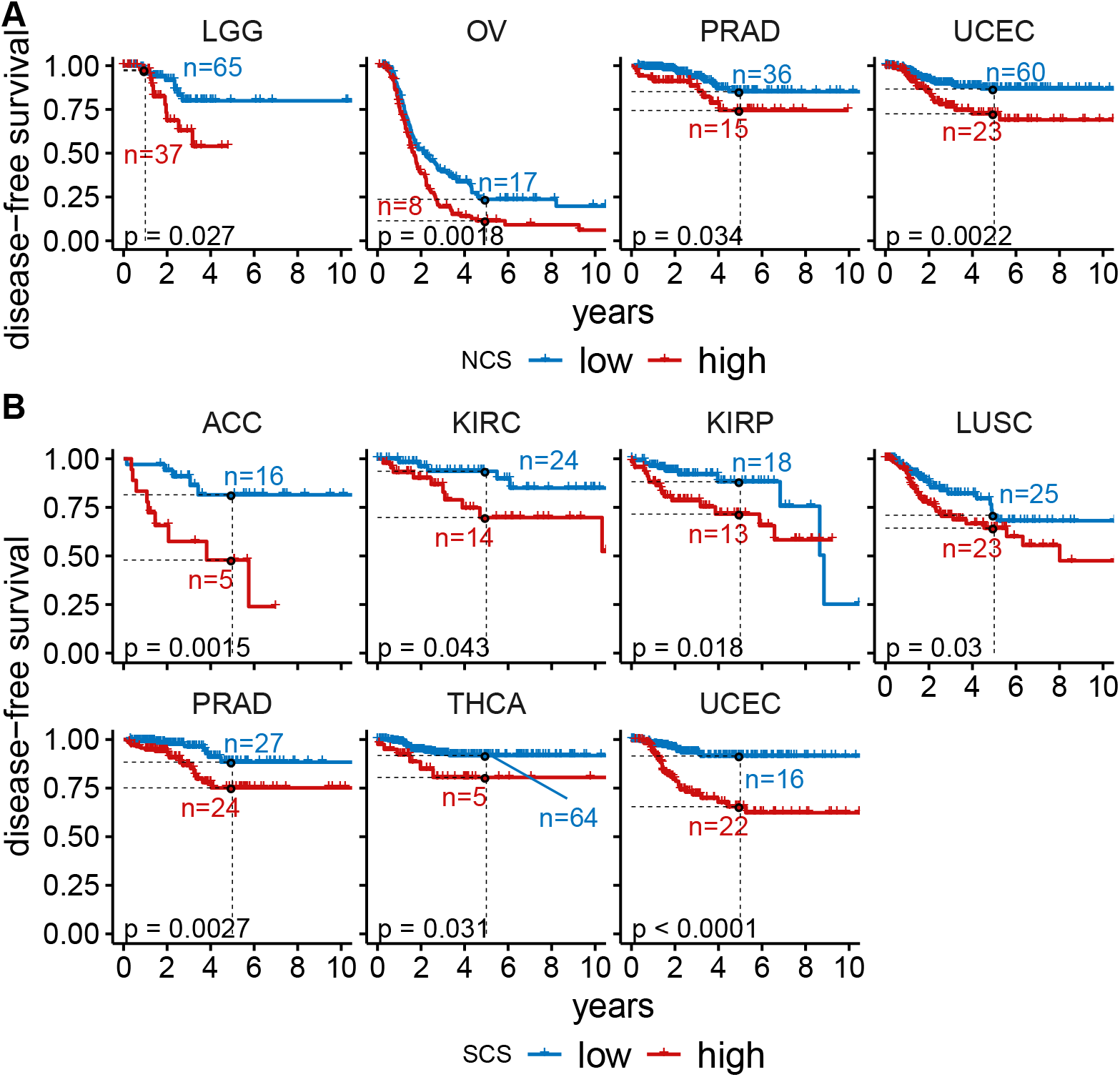
(**A**) Disease-free survival in four cancer types where significant differences between high- and low-NCS groups were observed. (**B**) Disease-free survival in seven cancer types where significant differences between high- and low-SCS group were observed.

**Figure A4.**
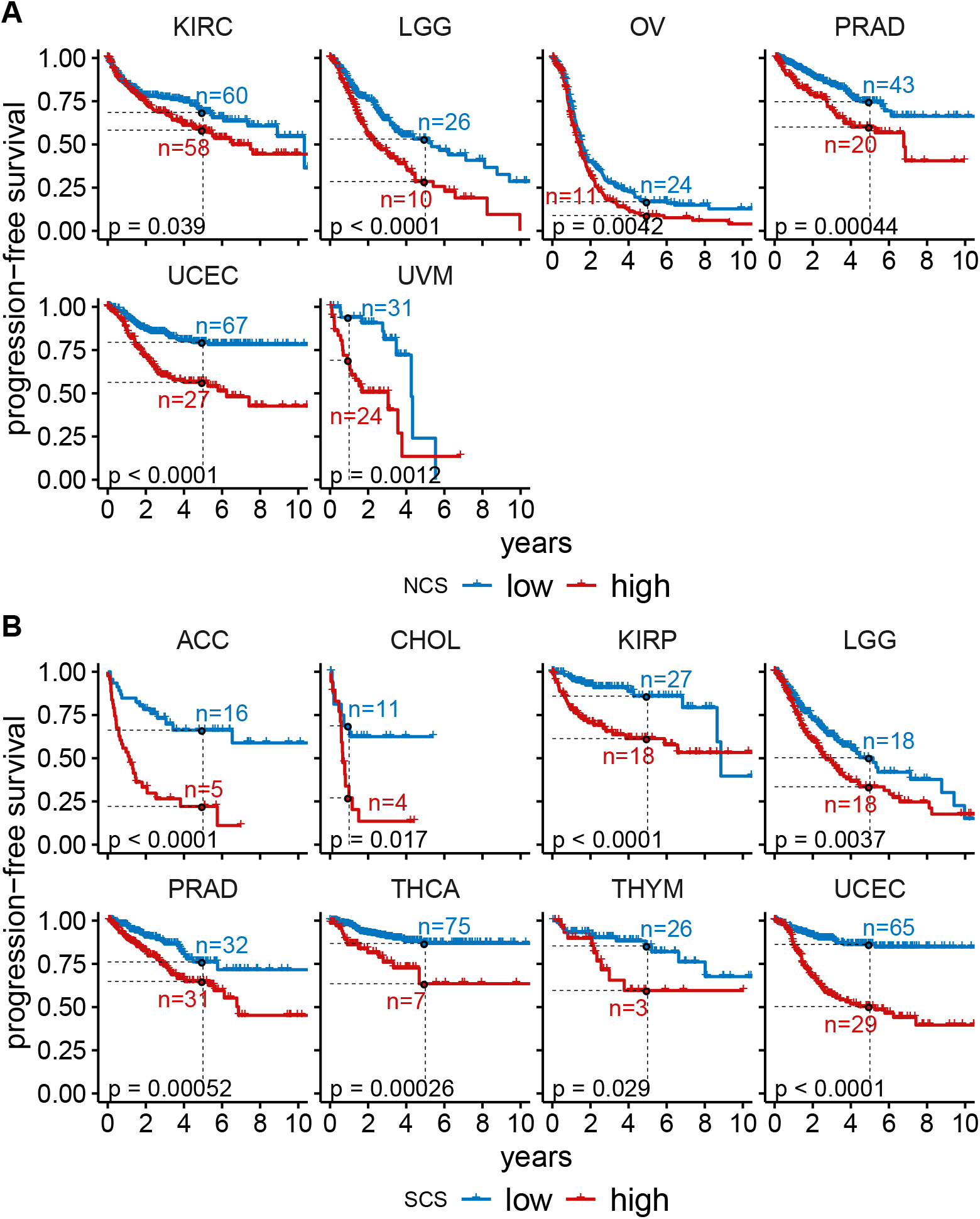
(**A**) Progression-free survival in six cancer types where significant differences between high- and low-NCS groups were observed. (**B**) Progression-free survival in eight cancer types where significant differences between high- and low-SCS groups were observed.

**Figure A5.**
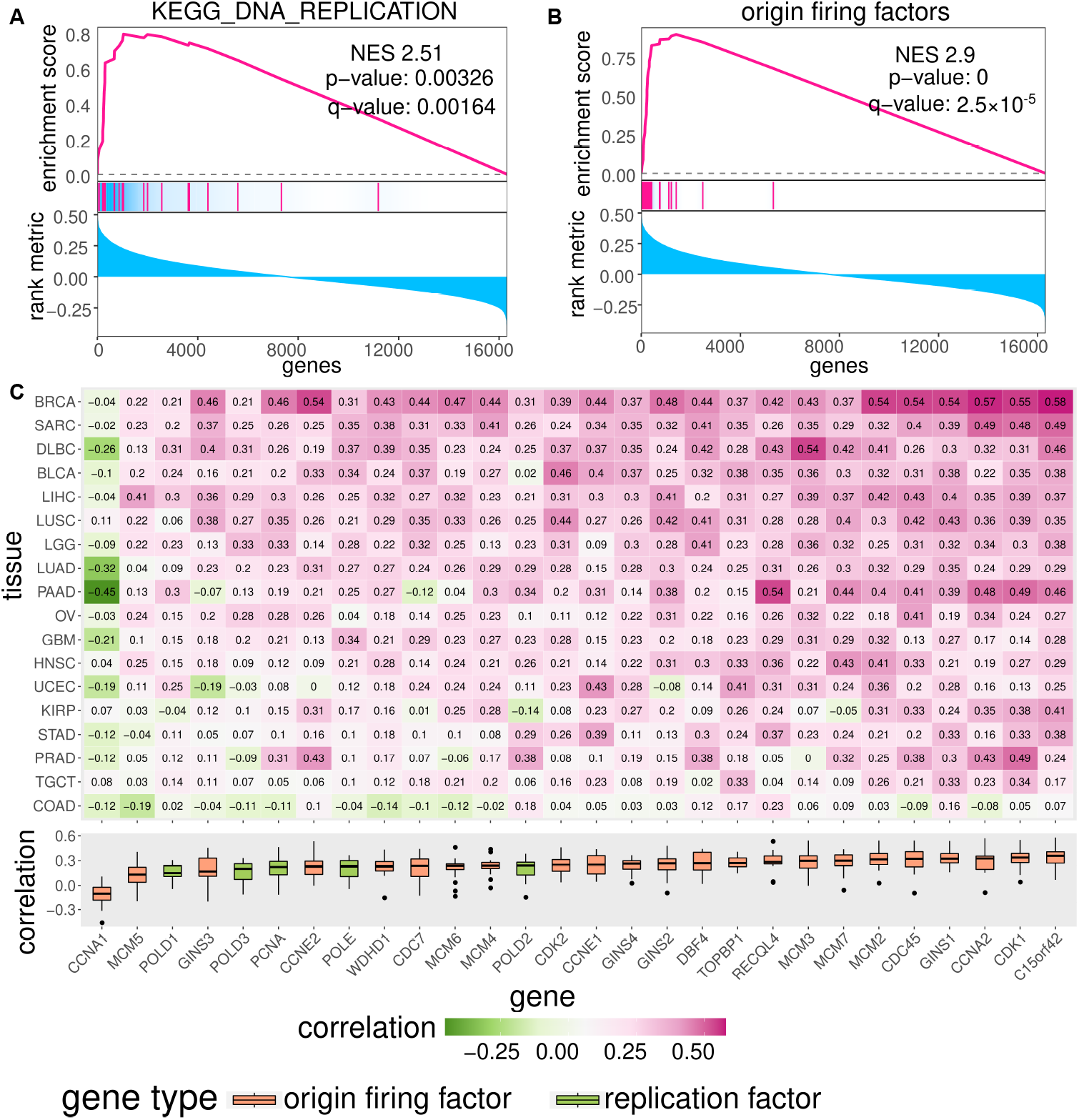
(**A**) Gene set enrichment analysis (GSEA) [67] for KEGG [68] DNA replication gene set. All genes are ordered according to the correlation of their expression with the SCS and enrichment significance is evaluated using permutation test. (**B**) GSEA analysis for manually curated origin firing factor gene set (curated in [10]). (**C**) Gene expression of many origin firing factors is positively correlated with SCS in many cancer types. Rows and columns of the heatmap represent cancer types and origin firing factor genes, respectively. Cancer types are clustered based on their correlation coefficients with origin firing factors. Genes are ordered based on the median correlation coefficient. Colour and values encoded in the heatmap represent the Spearman correlation coefficient.

**Figure A6.**
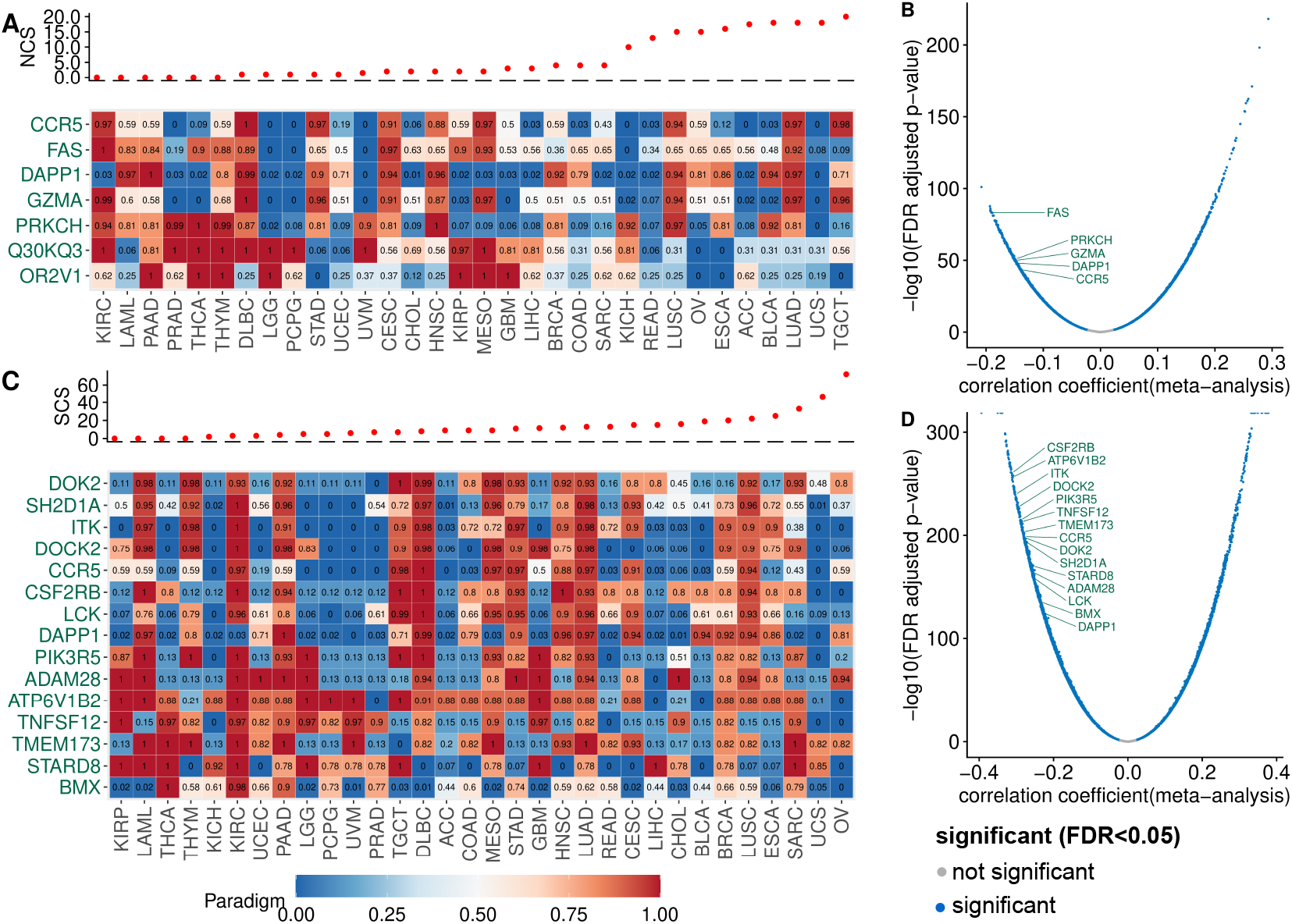
Paradigm pathway activity and gene expression negatively associated with CIN. (**A**) The PARADIGM pathway-level activities corresponding to protein-coding genes (rows) were correlated with the NCS. Only pathways with a significant negative correlation (FAD-adjusted *p <* 5%) less than *™*0.3 in at least seven cancer types were included. The heatmap shows the normalised PARADIGM pathway activity (0–1 from low to high). Cancer types are ordered according to their median NCS, see top panel. (**B**) Volcano plot for the correlation between gene expression and NCS, highlighting gene names corresponding to PARADIGM features that are significantly negatively associated with NCS. (**C**) Analogous to (**A**), but for the SCS instead of the NCS. (**D**) Correlation of SCS and gene expression, analogous to (**B**).

**Figure A7.**
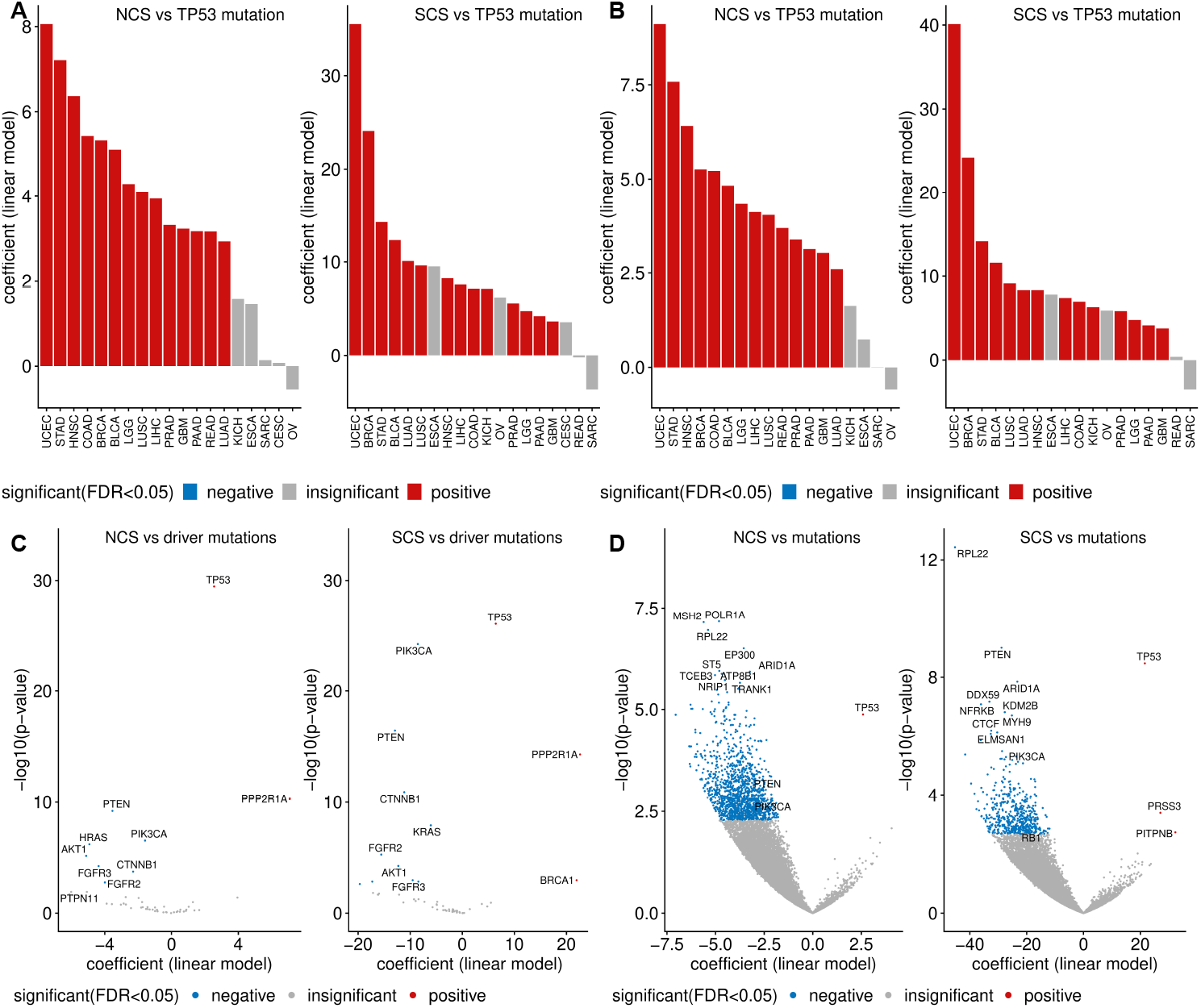
(**A**) *TP53* mutation is positively associated with high W-CIN in multiple cancer types. The bar shows the linear regression model coefficient using NCS as dependent variable and *TP53* mutation as explanatory variable. Only cancer types with both *TP53* mutant and *TP53* wild type in *≥*20 samples are considered. (**B**) *TP53* mutation is positively associated with SCS, the association analysis is performed as in (**A**), except excluding MIN samples. (**C**) The volcano plot shows the association between CIN and validated driver mutations, association analysis is performed using non-MIN samples only. (**D**) The volcano plot shows the correlation between CIN score and somatic mutations in CCLE cell line samples. For all associations in (**A**–**D**), red, grey and blue encode positive, insignificant and negative associations. FAD-adjusted *p ≤* 0.05 is considered as significant. Gene names of known important oncogenes and CIN driver genes are annotated in the volcano plot, if significantly associated with CIN.

## References

[1] Carter, S. L., Eklund, A. C., Kohane, I. S., Harris, L. N. & Szallasi, Z. A signature of chromosomal instability inferred from gene expression profiles predicts clinical outcome in multiple human cancers. Nat. Genet. 38, 1043–1048 (2006). URL http://dx.doi.org/10.1038/ng1861. https://doi.org/10.1038/ng1861.

[2] Bakhoum, S. F. & Cantley, L. C. The Multifaceted Role of Chromosomal Instability in Cancer and Its Microenvironment. Cell 174 (6), 1347–1360 (2018). URL https://linkinghub.elsevier.com/retrieve/pii/S0092867418310432. https://doi.org/10.1016/j.cell.2018.08.027.

[3] Sansregret, L., Vanhaesebroeck, B. & Swanton, C. Determinants and clinical implications of chromosomal instability in cancer. Nat Rev Clin Oncol 15 (3), 139–150 (2018). URL http://www.nature.com/articles/nrclinonc.2017.198. https://doi.org/10.1038/nrclinonc.2017.198.

[4] Roschke, A. & Rozenblum, E. Multi-layered cancer chromosomal instability phenotype. Front. Oncol. 3, 302 (2013). URL https://www.frontiersin.org/article/10.3389/fonc.2013.00302. https://doi.org/10.3389/fonc.2013.00302.

[5] Thompson, S. L., Bakhoum, S. F. & Compton, D. A. Mechanisms of chromosomal instability. Curr. Biol. 20, R285–R295 (2010). URL http://www.sciencedirect.com/science/article/pii/S096098221000076X. https://doi.org/10.1016/j.cub.2010.01.034.

[6] Siri, S. O., Martino, J. & Gottifredi, V. Structural Chromosome Instability: Types, Origins, Consequences, and Therapeutic Opportunities. Cancers 13 (12), 3056 (2021). URL https://www.mdpi.com/2072-6694/13/12/3056. https://doi.org/10.3390/cancers13123056.

[7] Burrell, R. A. et al. Replication stress links structural and numerical cancer chromosomal instability. Nature 494, 492–496 (2013). https://doi.org/10.1038/nature11935.

[8] Wilhelm, T., Said, M. & Naim, V. DNA Replication Stress and Chromosomal Instability: Dangerous Liaisons. Genes 11 (6), 642 (2020). URL https://www.mdpi.com/2073-4425/11/6/642. https://doi.org/10.3390/genes11060642.

[9] Böhly, N., Kistner, M. & Bastians, H. Mild replication stress causes aneuploidy by deregulating microtubule dynamics in mitosis. Cell Cycle 18 (20), 2770–2783 (2019). URL https://www.tandfonline.com/doi/full/10.1080/15384101.2019.1658477. https://doi.org/10.1080/15384101.2019.1658477.

[10] Schmidt, A.-K. et al. Dormant replication origin firing links replication stress to whole chromosomal instability in human cancer. bioRxiv (2021). URL http://biorxiv.org/lookup/doi/10.1101/2021.10.11.463929. https://doi.org/10.1101/2021.10.11.463929.

[11] Passerini, V. et al. The presence of extra chromosomes leads to genomic instability. Nat. Commun. 7 (1), 10754 (2016). https://doi.org/10.1038/ncomms10754.

[12] Dewhurst, S. M. et al. Tolerance of Whole-Genome Doubling Propagates Chromosomal Instability and Accelerates Cancer Genome Evolution. Cancer Discov 4 (2), 175–185 (2014). URL http://cancerdiscovery.aacrjournals.org/lookup/doi/10.1158/2159-8290.CD-13-0285. https://doi.org/10.1158/2159-8290.CD-13-0285.

[13] Bakhoum, S. F., Danilova, O. V., Kaur, P., Levy, N. B. & Compton, D. A. Chromosomal Instability Substantiates Poor Prognosis in Patients with Diffuse Large B-cell Lymphoma. Clin. Cancer Res. 17 (24), 7704–7711 (2011). URL http://clincancerres.aacrjournals.org/cgi/doi/10.1158/1078-0432.CCR-11-2049. https://doi.org/10.1158/1078-0432.CCR-11-2049.

[14] Tijhuis, A. E., Johnson, S. C. & McClelland, S. E. The emerging links between chromosomal instability (CIN), metastasis, inflammation and tumour immunity. Mol. Cytogenet. 12 (1), 17 (2019). URL https://doi.org/10.1186/s13039-019-0429-1. https://doi.org/10.1186/s13039-019-0429-1.

[15] Roylance, R. et al. Relationship of extreme chromosomal instability with long-term survival in a retrospective analysis of primary breast cancer. Cancer Epidemiol. Biomarkers Prev. 20 (10), 2183–2194 (2011). https://doi.org/10.1158/1055-9965.EPI-11-0343.

[16] Birkbak, N. J. et al. Paradoxical Relationship between Chromosomal Instability and Survival Outcome in Cancer. Cancer Res. 71 (10), 3447–3452 (2011). URL http://cancerres.aacrjournals.org/lookup/doi/10.1158/0008-5472.CAN-10-3667. https://doi.org/10.1158/0008-5472.CAN-10-3667.

[17] Gronroos, E. & López-García, C. Tolerance of Chromosomal Instability in Cancer: Mechanisms and Therapeutic Opportunities. Cancer Res. 78 (23), 6529–6535 (2018). URL http://cancerres.aacrjournals.org/lookup/doi/10.1158/0008-5472.CAN-18-1958. https://doi.org/10.1158/0008-5472.CAN-18-1958.

[18] Thompson, S. L. & Compton, D. A. Proliferation of aneuploid human cells is limited by a p53-dependent mechanism. J. Cell Biol. 188 (3), 369–381 (2010). URL https://rupress.org/jcb/article/188/3/369/35591/Proliferation-of-aneuploid-human-cells-is-limited. https://doi.org/10.1083/jcb.200905057.

[19] Sheltzer, J. M. A Transcriptional and Metabolic Signature of Primary Aneuploidy Is Present in Chromosomally Unstable Cancer Cells and Informs Clinical Prognosis. Cancer Res. 73 (21), 6401–6412 (2013). URL http://cancerres.aacrjournals.org/cgi/doi/10.1158/0008-5472.CAN-13-0749. https://doi.org/10.1158/0008-5472.CAN-13-0749.

[20] Endesfelder, D. et al. Chromosomal instability selects gene copy-number variants encoding core regulators of proliferation in er+ breast cancer. Cancer Res. 74 (17), 4853–4863 (2014). URL https://cancerres.aacrjournals.org/content/74/17/4853. https://doi.org/10.1158/0008-5472.CAN-13-2664.

[21] Buccitelli, C. et al. Pan-cancer analysis distinguishes transcriptional changes of aneuploidy from proliferation. Genome Res. 27 (4), 501–511 (2017). URL http://genome.cshlp.org/lookup/doi/10.1101/gr.212225.116. https://doi.org/10.1101/gr.212225.116.

[22] Davoli, T., Uno, H., Wooten, E. C. & Elledge, S. J. Tumor aneuploidy correlates with markers of immune evasion and with reduced response to immunotherapy. Science 355 (6322), eaaf8399 (2017). https://doi.org/10.1126/science.aaf8399.

[23] Bakhoum, S. F. et al. Chromosomal instability drives metastasis through a cytosolic DNA response. Nature 553, 467–472 (2018). URL https://doi.org/10.1038/nature25432. https://doi.org/10.1038/nature25432.

[24] Salgueiro, L. et al. Acquisition of chromosome instability is a mechanism to evade oncogene addiction. EMBO Mol Med 12 (3), e10941 (2020). URL https://onlinelibrary.wiley.com/doi/10.15252/emmm.201910941. https://doi.org/10.15252/emmm.201910941.

[25] Lukow, D. A. et al. Chromosomal instability accelerates the evolution of resistance to anti-cancer therapies. Dev. Cell 56 (17), 2427–2439.e4 (2021). URL https://linkinghub.elsevier.com/retrieve/pii/S153458072100592X. https://doi.org/10.1016/j.devcel.2021.07.009.

[26] Lee, A. J. et al. Chromosomal instability confers intrinsic multidrug resistance. Cancer Res. 71, 1858–1870 (2011). URL http://cancerres.aacrjournals.org/content/71/5/1858. https://doi.org/10.1158/0008-5472.CAN-10-3604.

[27] Taylor, A. M. et al. Genomic and Functional Approaches to Understanding Cancer Aneuploidy. Cancer Cell 33 (4), 676–689.e3 (2018). https://doi.org/10.1016/j.ccell.2018.03.007.

[28] Chang, K. et al. The Cancer Genome Atlas Pan-Cancer analysis project. Nat. Genet. 45 (10), 1113–1120 (2013). URL https://doi.org/10.1038/ng.2764. https://doi.org/10.1038/ng.2764.

[29] Carter, S. L. et al. Absolute quantification of somatic DNA alterations in human cancer. Nat. Biotechnol. 30 (5), 413–421 (2012). URL https://doi.org/10.1038/nbt.2203. https://doi.org/10.1038/nbt.2203.

[30] Vaske, C. J. et al. Inference of patient-specific pathway activities from multidimensional cancer genomics data using PARADIGM. Bioinformatics 26 (12), i237–i245 (2010). URL https://doi.org/10.1093/bioinformatics/btq182. https://doi.org/10.1093/bioinformatics/btq182.

[31] Barretina, J. et al. The Cancer Cell Line Encyclopedia enables predictive modelling of anticancer drug sensitivity. Nature 483 (7391), 603–607 (2012). URL https://doi.org/10.1038/nature11003. https://doi.org/10.1038/nature11003.

[32] Seashore-Ludlow, B. et al. Harnessing connectivity in a large-scale small-molecule sensitivity dataset. Cancer Discov 5 (11), 1210–1223 (2015). URL https://cancerdiscovery.aacrjournals.org/content/5/11/1210. https://doi.org/10.1158/2159-8290.CD-15-0235.

[33] Thorsson, V. et al. The immune landscape of cancer. Immunity 48 (4), 812–830.e14 (2018). URL http://www.sciencedirect.com/science/article/pii/S1074761318301213. https://doi.org/10.1016/j.immuni.2018.03.023.

[34] Bailey, M. H. et al. Comprehensive Characterization of Cancer Driver Genes and Mutations. Cell 173 (2), 371–385.e18 (2018). https://doi.org/10.1016/j.cell.2018.02.060.

[35] Therneau, T. M. A Package for Survival Analysis in R. R package version 3.2-10 (2021). Available online: https://CRAN.R-project.org/package=survival (accessed on 4 October 2021).

[36] Kassambara, A., Kosinski, M. & Biecek, P. survminer: Drawing Survival Curves using ‘ggplot2’. R package version 0.4.9 (2021). Available online: https://CRAN.R-project.org/package=survminer (accessed on 4 October2021).

[37] Ritchie, M. E. et al. limma powers differential expression analyses for RNA-sequencing and microarray studies. Nucleic Acids Res. 43 (7), e47 (2015). https://doi.org/10.1093/nar/gkv007.

[38] Lengauer, C., Kinzler, K. W. & Vogelstein, B. Genetic instabilities in human cancers. Nature 396 (6712), 643–649 (1998). URL https://doi.org/10.1038/25292. https://doi.org/10.1038/25292.

[39] McGranahan, N., Burrell, R. A., Endesfelder, D., Novelli, M. R. & Swanton, C. Cancer chromosomal instability: Therapeutic and diagnostic challenges. EMBO Rep. 13 (6), 528–538 (2012). https://doi.org/10.1038/embor.2012.61.

[40] Lepage, C. C., Morden, C. R., Palmer, M. C. L., Nachtigal, M. W. & McManus, K. J. Detecting chromosome instability in cancer: Approaches to resolve cell-to-cell heterogeneity. Cancers 11 (2), 226 (2019). URL https://pubmed.ncbi.nlm.nih.gov/30781398. https://doi.org/10.3390/cancers11020226.

[41] Delaney, J. R. et al. Haploinsufficiency networks identify targetable patterns of allelic deficiency in low mutation ovarian cancer. Nat. Commun. 8 (1), 14423 (2017). URL https://doi.org/10.1038/ncomms14423. https://doi.org/10.1038/ncomms14423.

[42] van Jaarsveld, R. H. & Kops, G. J. Difference makers: Chromosomal instability versus aneuploidy in cancer. Trends Cancer 2 (10), 561–571 (2016). URL https://www.sciencedirect.com/science/article/pii/S2405803316301200. https://doi.org/10.1016/j.trecan.2016.09.003.

[43] Sheltzer, J. M. & Amon, A. The aneuploidy paradox: costs and benefits of an incorrect karyotype. Trends Genet. 27 (11), 446–453 (2011). URL https://linkinghub.elsevier.com/retrieve/pii/S0168952511001181. https://doi.org/10.1016/j.tig.2011.07.003.

[44] Salmina, K. et al. The Cancer Aneuploidy Paradox: In the Light of Evolution. Genes 10 (2), 83 (2019). https://doi.org/10.3390/genes10020083.

[45] Chunduri, N. K. & Storchová, Z. The diverse consequences of aneuploidy. Nat. Cell Biol. 21 (1), 54–62 (2019). URL http://www.nature.com/articles/s41556-018-0243-8. https://doi.org/10.1038/s41556-018-0243-8.

[46] Lord, C. J. & Ashworth, A. BRCAness revisited. Nat. Rev. Cancer 16 (2), 110–120 (2016). URL https://doi.org/10.1038/nrc.2015.21. https://doi.org/10.1038/nrc.2015.21.

[47] Turner, N., Tutt, A. & Ashworth, A. Hallmarks of ‘BRCAness’ in sporadic cancers. Nat. Rev. Cancer 4 (10), 814–819 (2004). URL https://doi.org/10.1038/nrc1457. https://doi.org/10.1038/nrc1457.

[48] Thompson, L., Jeusset, L., Lepage, C. & McManus, K. Evolving Therapeutic Strategies to Exploit Chromosome Instability in Cancer. Cancers 9 (12), 151 (2017). URL http://www.mdpi.com/2072-6694/9/11/151. https://doi.org/10.3390/cancers9110151. Numerical and structural chromosome instability across different cancer types 41

[49] Wu, Y.-L. et al. Afatinib versus cisplatin plus gemcitabine for first-line treatment of Asian patients with advanced non-small-cell lung cancer harbouring EGFR mutations (LUX-Lung 6): an open-label, randomised phase 3 trial. Lancet Oncol. 15 (2), 213–222 (2014). URL https://doi.org/10.1016/S1470-2045(13)70604-1. https://doi.org/10.1016/S1470-2045(13)70604-1.

[50] Bugler, B., Schmitt, E., Aressy, B. & Ducommun, B. Unscheduled expression of CDC25B in S-phase leads to replicative stress and DNA damage. Mol. Cancer 9 (1), 29 (2010). URL https://doi.org/10.1186/1476-4598-9-29. https://doi.org/10.1186/1476-4598-9-29.

[51] Kamada, K. in The GINS Complex: Structure and Function (ed.MacNeill, S.) The Eukaryotic Replisome: a Guide to Protein Structure and Function, Vol. 62 135–156 (Springer, Dordrecht, The Netherlands, 2012). URL http://link.springer.com/10.1007/978-94-007-4572-88.

[52] Shi, W. et al. CKS1B as Drug Resistance-Inducing Gene—A Potential Target to Improve Cancer Therapy. Front. Oncol. 10, 582451 (2020). URL https://www.frontiersin.org/article/10.3389/fonc.2020.582451/full. https://doi.org/10.3389/fonc.2020.582451.

[53] Eischen, C. M. Genome stability requires p53. Cold Spring Harb. Perspect. Med. 6 (6), a026096 (2016). URL http://perspectivesinmedicine.cshlp.org/content/6/6/a026096.abstract. https://doi.org/10.1101/cshperspect.a026096.

[54] Ciriello, G. et al. Emerging landscape of oncogenic signatures across human cancers. Nat. Genet. 45 (10), 1127–1133 (2013). URL http://www.nature.com/articles/ng.2762. https://doi.org/10.1038/ng.2762.

[55] Tamborero, D. et al. Cancer Genome Interpreter annotates the biological and clinical relevance of tumor alterations. Genome Med. 10 (1), 25 (2018). URL https://doi.org/10.1186/s13073-018-0531-8. https://doi.org/10.1186/s13073-018-0531-8.

[56] Carramusa, L. et al. The PVT-1 oncogene is a Myc protein target that is overexpressed in transformed cells. J. Cell. Physiol. 213 (2), 511–518 (2007). URL https://onlinelibrary.wiley.com/doi/10.1002/jcp.21133. https://doi.org/10.1002/jcp.21133.

[57] Zhang, C., Hu, H., Wang, X., Zhu, Y. & Jiang, M. Wfdc protein: A promising diagnosis biomarker of ovarian cancer. J. Cancer 12, 5404–5412 (2021). URL https://www.jcancer.org/v12p5404.htm. https://doi.org/10.7150/jca.57880.42 Numerical and structural chromosome instability across different cancer types

[58] L’ opez, S. et al. Interplay between whole-genome doubling and the accumulation of deleterious alterations in cancer evolution. Nat. Genet. 52 (3), 283–293 (2020). URL https://doi.org/10.1038/s41588-020-0584-7. https://doi.org/10.1038/s41588-020-0584-7.

[59] Quinton, R. J. et al. Whole-genome doubling confers unique genetic vulnerabilities on tumour cells. Nature 590 (7846), 492–497 (2021). URL https://www.nature.com/articles/s41586-020-03133-3. https://doi.org/10.1038/s41586-020-03133-3.

[60] Bakhoum, S. F., Kabeche, L., Murnane, J. P., Zaki, B. I. & Compton, D. A. DNA-Damage Response during Mitosis Induces Whole-Chromosome Missegregation. Cancer Discov 4 (11), 1281–1289 (2014). https://doi.org/10.1158/2159-8290.CD-14-0403.

[61] Bakhoum, S. F. et al. Numerical chromosomal instability mediates susceptibility to radiation treatment. Nat. Commun. 6 (1), 5990 (2015). URL http://www.nature.com/articles/ncomms6990. https://doi.org/10.1038/ncomms6990.

[62] Davoli, T. et al. Cumulative Haploinsufficiency and Triplosensitivity Drive Aneuploidy Patterns and Shape the Cancer Genome. Cell 155 (4), 948–962 (2013). URL https://linkinghub.elsevier.com/retrieve/pii/S0092867413012877. https://doi.org/10.1016/j.cell.2013.10.011.

[63] Berenjeno, I. M. et al. Oncogenic PIK3CA induces centrosome amplification and tolerance to genome doubling. Nat. Commun. 8 (1), 1773 (2017). URL http://www.nature.com/articles/s41467-017-02002-4. https://doi.org/10.1038/s41467-017-02002-4.

[64] Laughney, A., Elizalde, S., Genovese, G. & Bakhoum, S. Dynamics of Tumor Heterogeneity Derived from Clonal Karyotypic Evolution. Cell Rep. 12 (5), 809–820 (2015). URL https://linkinghub.elsevier.com/retrieve/pii/S2211124715007032. https://doi.org/10.1016/j.celrep.2015.06.065.

[65] Heng, H. et al. Karyotype Heterogeneity and Unclassified Chromosomal Abnormalities. Cytogenet. Genome Res. 139 (3), 144–157 (2013). URL https://www.karger.com/DOI/10.1159/000348682. https://doi.org/10.1159/000348682.

[66] Ye, C. J., Stilgenbauer, L., Moy, A., Liu, G. & Heng, H. H. What Is Karyotype Coding and Why Is Genomic Topology Important for Cancer and Evolution? Front. Genet. 10, 1082 (2019). https://doi.org/10.3389/fgene.2019.01082.

[67] Subramanian, A. et al. Gene set enrichment analysis: A knowledge-based approach for interpreting genome-wide expression profiles. Proc. Natl. Acad. Sci. U.S.A. 102 (43), 15545–15550 (2005). URL https://www.pnas.org/content/102/43/15545. https://doi.org/10.1073/pnas.0506580102.

[68] Kanehisa, M. & Goto, S. KEGG: Kyoto Encyclopedia of Genes and Genomes. Nucleic Acids Res. 28 (1), 27–30 (2000). URL https://doi.org/10.1093/nar/28.1.27. https://doi.org/10.1093/nar/28.1.27.

