## Supplemental Text for "Distinct and common features of numerical and structural chromosomal instability across different cancer types"

This PDF file includes pan-cancer weighted genome instability index (WGII) association analysis results. We performed the same association analysis on WGII as did on NCS/SCS in the main text.

### 1. Landscape of WGII

Consistent with results for NCS analysis in main text. Two clearly separated clusters of WGII are formed per cancer. The clusters are strongly associated with whole genome doubling (WGD) (Figure 1A). WGII is strongly and consistently associated with aneuploidy score as shown in Figure 1B. While WGII can distinguish metastatic tumours from primary tumours and normal tissues, NCS only differentiates normal tissue and primary tumours (Figure 2B).

Springer Nature 2021 LATEX template



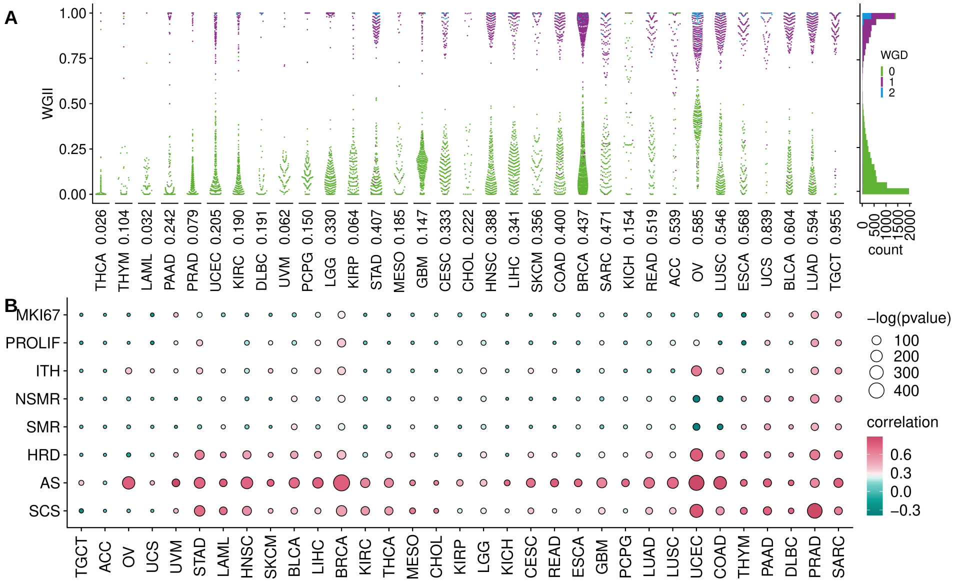


**Figure 1.** Distribution of WGII and their association with genetic instability. (**A**) Left: WGII for TCGA tumour samples (dots) from different cancer types, sorted according to median WGII. The colour coding indicates the WGD status and the number below each beeswarm plot is the proportion of samples which underwent WGD. Right: Pan-cancer histogram of WGII. (**B**) Correlation between WGII with different indices for genetic instability, intra-tumour heterogeneity and proliferation: *MKI67* expression, proliferation rates (PROLIF), intra-tumour heterogeneity (ITH), non-silent mutation rate (NSMR), silent mutation rate (SMR), homologous recombination deficiency (HRD) and aneuploidy score (AS), structural complexity score (SCS).

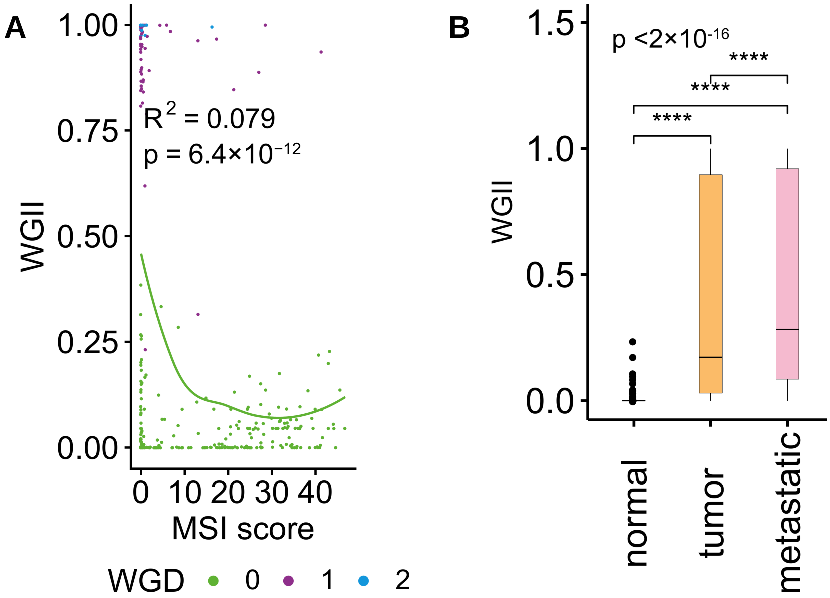


**Figure 2.** (**A**) The relationship between microsatellite instability (MIN) scores and the WGII. (**B**) Comparison of the WGII between normal samples with primary and metastatic tumour samples.

### 2. Clinical significance of WGII

Clinical associations of WGII show the same patterns as those performed using NCS (Figure 3), but WGII has better differentiation ability compared with NCS. Additional detailed clinical association analysis is in Section 6.

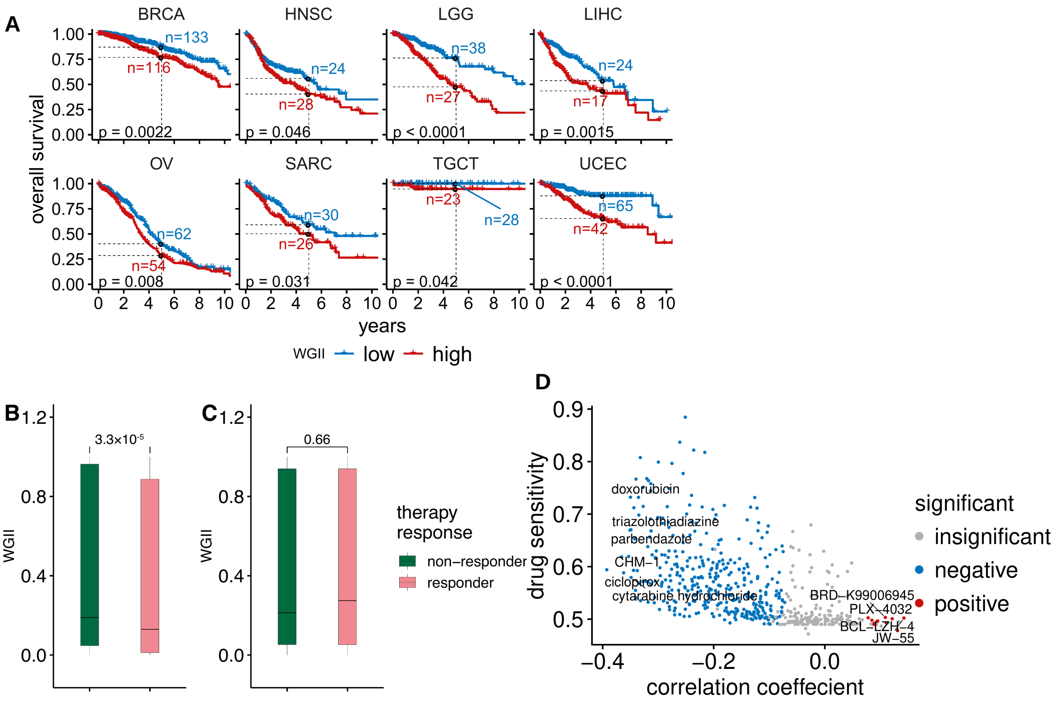


**Figure 3.** Clinical significance of WGII in different cancer types. (**A**) For eight cancer types there are significant differences in overall survival between patient samples with low WGII (blue) and high WGII (red). Dashed lines indicate the five year overall survival probability of the two groups. (**B**) Left: Comparison of the WGII between radiotherapy responders and non-responders using a Wilcoxon rank sum test. Right: WGII between chemotherapy responders and non-responders using a Wilcoxon rank sum test. **(C)** The median drug sensitivity of a compound plotted against the correlation coefficient between drug sensitivity and WGII. Drugs with significant positive and negative correlations between their sensitivity and WGII are highlighted in red and blue, respectively.

### 3. PARADIGM pathway activity and WGII

PARADIGM pathway activity associations of WGII show the same patterns as those performed using NCS (Figure 4), but WGII has better differentiation ability compared with NCS. Only 11 PARADIGM protein-coding genes are consistently and positively associated with NCS. More than 15 PARADIGM protein-coding genes are consistently and positively associated with WGII.

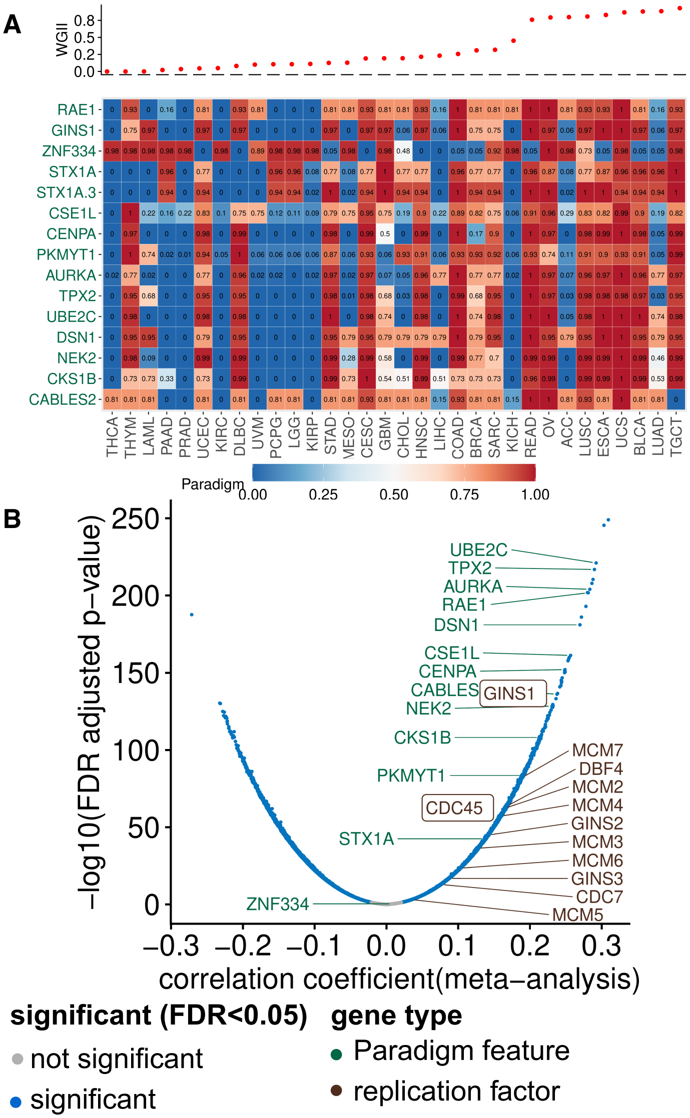


**Figure 4.** PARADIGM pathway activity and gene expression associated with WGII. (**A**) The PARADIGM pathway-level activities corresponding to protein-coding genes (rows) were correlated with the WGII. Only pathways with a significant correlation (FDR-adjusted $p < 5\%$) larger than 0.3 in at least seven cancer types were included. The heatmap shows the normalised PARADIGM pathway activity (0–1 from low to high). Cancer types were ordered according to their median WGII, see top panel. (**B**) Volcano plot for the correlation between gene expression and WGII.

### 4. Somatic point mutations and WGII

Somatic mutation associations of WGII show the same patterns as those performed using NCS (Figure 5).

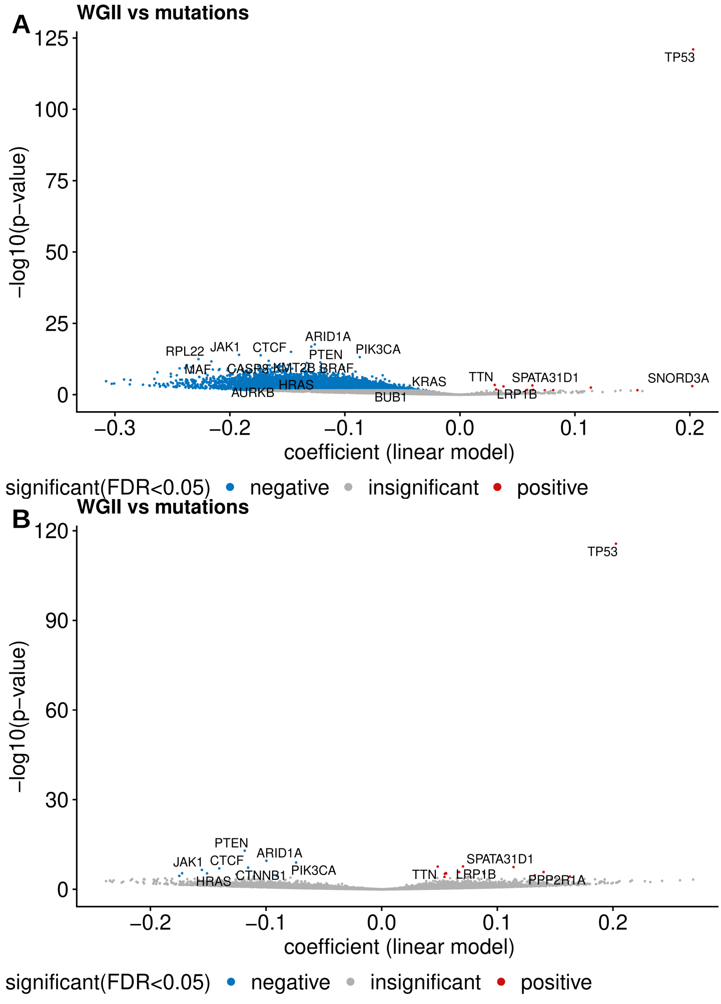


**Figure 5.** Pan-cancer somatic mutations and WGII. (**A**) The volcano plots show the association between somatic mutations and the WGII. The linear model coefficient indicates the mean difference of the respective WGII when the mutation is present in a tumour sample relative to the wild type. Genes with lowest p-values, well-known CIN genes and cancer driver genes are highlighted. The analysis was performed on genes for which samples sizes for both wild type group and mutated group are larger than 19. Mutations significantly associated (FDR-adjusted $p < 5\%$) with higher or lower WGII are highlighted in blue and red, respectively. (**B**) The same as (**A**), but hypermutated MIN samples are excluded.

### 5. Copy number changes and WGII

Copy number associations of WGII show the same patterns as those performed using NCS (Figure 6).

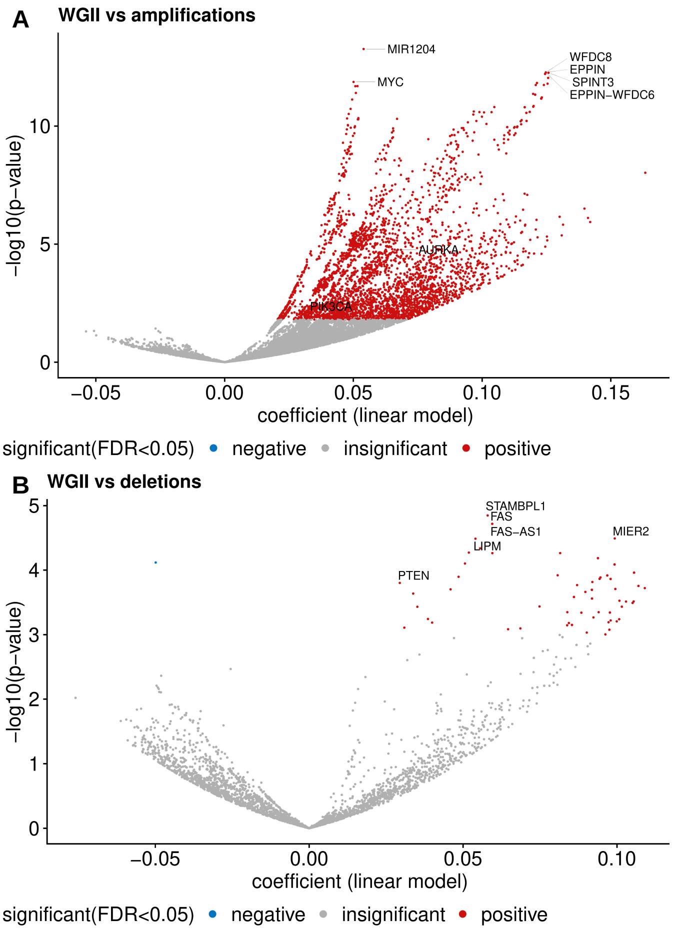


**Figure 6.** Copy number amplifications and deletions enriched in high WGII samples. (**A**) The volcano plots show the gene wise associations between copy number amplification status and WGII, obtained from a regression model adjusted by cancer type. The linear model coefficient indicates the mean difference of the respective WGII when the alteration is present in a tumour sample relative to the wild type. Genes with the lowest p-values and well-known CIN genes are highlighted. Blue and red colours encode genes with a significantly higher alteration frequency (FDR-adjusted $p < 5\%$) in samples with low and high WGII, respectively. The analysis was performed on 16,922 genes with sample sizes greater than 19 for both wild type and amplified groups. (**B**) Pan-cancer copy number deletions associated with WGII displayed in an analogous way as in (**A**).

### 6. Additional clinical association analysis

**Table 1.** Association between WGII and overall survival across cancer types

| cohort | sample_number | pvalue | low_surv5[a] | high_surv5[b] | low_surv5_n[c] | high_surv5_n[d] |
|---|---|---|---|---|---|---|
| LGG | 509 | 0.00 | 0.76 | 0.48 | 38.00 | 27.00 |
| UCEC | 518 | 0.00 | 0.88 | 0.65 | 65.00 | 42.00 |
| LIHC | 366 | 0.00 | 0.54 | 0.44 | 24.00 | 17.00 |
| BRCA | 1066 | 0.00 | 0.87 | 0.77 | 133.00 | 116.00 |
| OV | 558 | 0.01 | 0.40 | 0.29 | 62.00 | 54.00 |
| SARC | 252 | 0.03 | 0.59 | 0.50 | 30.00 | 26.00 |
| TGCT | 133 | 0.04 | 1.00 | 0.95 | 28.00 | 23.00 |
| HNSC | 516 | 0.05 | 0.56 | 0.40 | 24.00 | 28.00 |
| UVM* | 80 | 0.07 | 0.94 | 0.92 | 33.00 | 31.00 |
| THCA | 497 | 0.09 | 0.97 | 0.89 | 51.00 | 45.00 |
| THYM | 122 | 0.09 | 0.98 | 0.88 | 18.00 | 15.00 |
| KIRP | 282 | 0.10 | 0.84 | 0.68 | 34.00 | 19.00 |
| PCPG | 161 | 0.14 | 0.98 | 0.95 | 16.00 | 12.00 |
| GBM | 571 | 0.17 | 0.07 | 0.06 | 10.00 | 9.00 |
| ACC | 89 | 0.18 | 0.69 | 0.56 | 14.00 | 14.00 |
| DLBC | 48 | 0.19 | 0.72 | 0.90 | 6.00 | 3.00 |
| STAD | 433 | 0.25 | 0.41 | 0.35 | 10.00 | 8.00 |
| CESC | 294 | 0.27 | 0.73 | 0.60 | 25.00 | 16.00 |
| LAML | 179 | 0.34 | 0.24 | 0.17 | 6.00 | 3.00 |
| PRAD | 489 | 0.41 | 0.99 | 0.97 | 35.00 | 49.00 |
| KIRC | 506 | 0.44 | 0.62 | 0.63 | 66.00 | 81.00 |
| CHOL* | 36 | 0.45 | 0.71 | 0.88 | 12.00 | 15.00 |
| READ* | 154 | 0.48 | 0.94 | 0.96 | 51.00 | 64.00 |
| ESCA* | 182 | 0.49 | 0.82 | 0.70 | 65.00 | 50.00 |
| MESO* | 86 | 0.59 | 0.65 | 0.71 | 28.00 | 28.00 |
| LUAD | 491 | 0.63 | 0.42 | 0.40 | 28.00 | 25.00 |
| COAD | 425 | 0.75 | 0.66 | 0.54 | 25.00 | 15.00 |
| KICH | 65 | 0.77 | 0.86 | 0.85 | 17.00 | 20.00 |
| SKCM* | 104 | 0.77 | 0.84 | 0.91 | 33.00 | 37.00 |
| PAAD | 183 | 0.85 | 0.16 | 0.33 | 3.00 | 5.00 |
| BLCA | 405 | 0.87 | 0.42 | 0.42 | 24.00 | 23.00 |
| UCS* | 56 | 0.88 | 0.79 | 0.81 | 22.00 | 20.00 |
| LUSC | 481 | 0.96 | 0.50 | 0.45 | 38.00 | 42.00 |

[a] 5-year overall survival probability in low WGII group.
[b] 5-year overall survival probability in high WGII group.
[c] number of samples at risk in low WGII group at 5th year.
[d] number of samples at risk in high WGII group at 5th year.
[*] 1-year overall survival statistics was reported in these cancer types due to short survival.

**Table 2.** Association between WGII and disease free survival across cancer types

| **cohort** | **sample_number** | **pvalue** | **low_surv5**[a] | **high_surv5**[b] | **low_surv5_n**[c] | **high_surv5_n**[d] |
|---|---|---|---|---|---|---|
| UCEC | 406 | 0.00 | 0.90 | 0.72 | 55.00 | 28.00 |
| OV | 279 | 0.00 | 0.25 | 0.10 | 17.00 | 8.00 |
| PRAD | 332 | 0.02 | 0.87 | 0.78 | 24.00 | 27.00 |
| LIHC | 315 | 0.03 | 0.41 | 0.22 | 14.00 | 4.00 |
| COAD | 175 | 0.05 | 0.83 | 0.62 | 11.00 | 3.00 |
| LGG* | 130 | 0.06 | 0.97 | 0.98 | 61.00 | 41.00 |
| LUSC | 295 | 0.08 | 0.73 | 0.63 | 24.00 | 24.00 |
| MESO* | 15 | 0.08 | 0.67 | 1.00 | 6.00 | 2.00 |
| CESC | 170 | 0.11 | 0.85 | 0.76 | 17.00 | 9.00 |
| KICH | 29 | 0.16 | 0.91 | 1.00 | 5.00 | 12.00 |
| BRCA | 927 | 0.25 | 0.86 | 0.82 | 105.00 | 85.00 |
| CHOL* | 24 | 0.26 | 0.71 | 0.50 | 10.00 | 4.00 |
| DLBC | 28 | 0.28 | 1.00 | 0.92 | 5.00 | 3.00 |
| UCS* | 26 | 0.28 | 1.00 | 0.77 | 11.00 | 9.00 |
| PAAD* | 68 | 0.35 | 0.86 | 0.80 | 21.00 | 21.00 |
| KIRC | 107 | 0.38 | 0.92 | 0.76 | 19.00 | 19.00 |
| TGCT | 104 | 0.39 | 0.68 | 0.82 | 9.00 | 15.00 |
| SARC | 148 | 0.44 | 0.56 | 0.45 | 16.00 | 11.00 |
| GBM* | 3 | 0.48 | 1.00 | 1.00 | 1.00 | 2.00 |
| KIRP | 180 | 0.52 | 0.74 | 0.89 | 17.00 | 14.00 |
| READ* | 42 | 0.53 | 0.90 | 1.00 | 15.00 | 20.00 |
| LUAD | 291 | 0.59 | 0.63 | 0.54 | 19.00 | 18.00 |
| THCA | 352 | 0.76 | 0.91 | 0.90 | 36.00 | 33.00 |
| ESCA* | 87 | 0.80 | 0.75 | 0.82 | 28.00 | 22.00 |
| ACC | 52 | 0.83 | 0.68 | 0.72 | 11.00 | 10.00 |
| BLCA | 187 | 0.88 | 0.72 | 0.71 | 11.00 | 15.00 |
| HNSC | 130 | 0.94 | 0.69 | 0.54 | 7.00 | 4.00 |
| STAD | 255 | 0.96 | 0.59 | 0.70 | 7.00 | 6.00 |
| PCPG | 144 | 0.97 | 0.95 | 0.97 | 12.00 | 10.00 |

[a] 5-year disease free survival probability in low WGII group.
[b] 5-year disease free survival probability in high WGII group.
[c] number of samples at risk in low WGII group at 5th year.
[d] number of samples at risk in high WGII group at 5th year.
[*] 1-year disease-free survival statistics was reported in these cancer types due to short survival.

**Table 3.** Association between WGII and progression free survival across cancer types

| cohort | sample_number | pvalue | low_surv5[a] | high_surv5[b] | low_surv5_n[c] | high_surv5_n[d] |
|---|---|---|---|---|---|---|
| UCEC | 518 | 0.00 | 0.85 | 0.56 | 60.00 | 34.00 |
| LGG | 509 | 0.00 | 0.55 | 0.28 | 26.00 | 10.00 |
| PRAD | 489 | 0.00 | 0.75 | 0.66 | 27.00 | 36.00 |
| OV | 558 | 0.00 | 0.18 | 0.08 | 24.00 | 11.00 |
| LIHC | 366 | 0.00 | 0.35 | 0.19 | 13.00 | 5.00 |
| UVM* | 79 | 0.01 | 0.94 | 0.69 | 31.00 | 24.00 |
| THYM | 122 | 0.01 | 0.91 | 0.65 | 17.00 | 12.00 |
| SKCM* | 104 | 0.03 | 0.78 | 0.64 | 28.00 | 24.00 |
| CHOL* | 36 | 0.06 | 0.63 | 0.34 | 10.00 | 5.00 |
| KIRC | 504 | 0.06 | 0.68 | 0.59 | 55.00 | 63.00 |
| CESC | 294 | 0.06 | 0.71 | 0.62 | 22.00 | 14.00 |
| ESCA* | 182 | 0.07 | 0.66 | 0.58 | 47.00 | 37.00 |
| ACC | 89 | 0.08 | 0.50 | 0.39 | 11.00 | 10.00 |
| SARC | 252 | 0.14 | 0.44 | 0.34 | 20.00 | 15.00 |
| HNSC | 516 | 0.20 | 0.51 | 0.46 | 20.00 | 24.00 |
| BRCA | 1066 | 0.22 | 0.80 | 0.76 | 119.00 | 102.00 |
| BLCA | 406 | 0.29 | 0.38 | 0.44 | 19.00 | 17.00 |
| READ* | 154 | 0.32 | 0.90 | 0.88 | 47.00 | 57.00 |
| GBM | 571 | 0.33 | 0.04 | 0.02 | 5.00 | 3.00 |
| TGCT | 133 | 0.43 | 0.69 | 0.80 | 17.00 | 20.00 |
| COAD | 425 | 0.43 | 0.63 | 0.55 | 21.00 | 9.00 |
| DLBC | 48 | 0.43 | 0.70 | 0.77 | 6.00 | 3.00 |
| KIRP | 281 | 0.48 | 0.72 | 0.79 | 28.00 | 17.00 |
| LUSC | 482 | 0.53 | 0.55 | 0.54 | 31.00 | 34.00 |
| LUAD | 491 | 0.68 | 0.39 | 0.39 | 20.00 | 18.00 |
| THCA | 497 | 0.74 | 0.85 | 0.83 | 42.00 | 40.00 |
| KICH | 65 | 0.75 | 0.87 | 0.87 | 17.00 | 20.00 |
| STAD | 435 | 0.82 | 0.40 | 0.48 | 10.00 | 8.00 |
| PCPG | 161 | 0.83 | 0.80 | 0.87 | 11.00 | 11.00 |
| PAAD* | 183 | 0.85 | 0.62 | 0.64 | 42.00 | 50.00 |
| UCS* | 56 | 0.92 | 0.50 | 0.58 | 14.00 | 14.00 |
| MESO* | 84 | 0.97 | 0.59 | 0.52 | 22.00 | 19.00 |

[a] 5-year progression free survival probability in low WGII group.
[b] 5-year progression free survival probability in high WGII group.
[c] number of samples at risk in low WGII group at 5th year.
[d] number of samples at risk in high WGII group at 5th year.
[*] 1-year progression-free survival statistics was reported in these cancer types due to short survival.

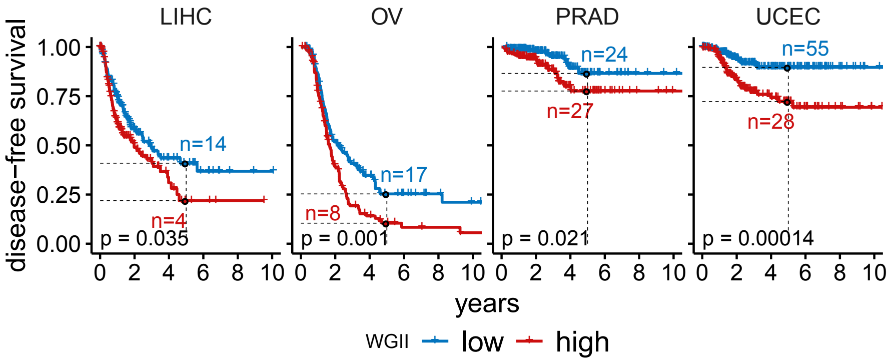


**Figure 7.** Disease free survival in four cancer types where significant differences between high and low WGII group were observed.

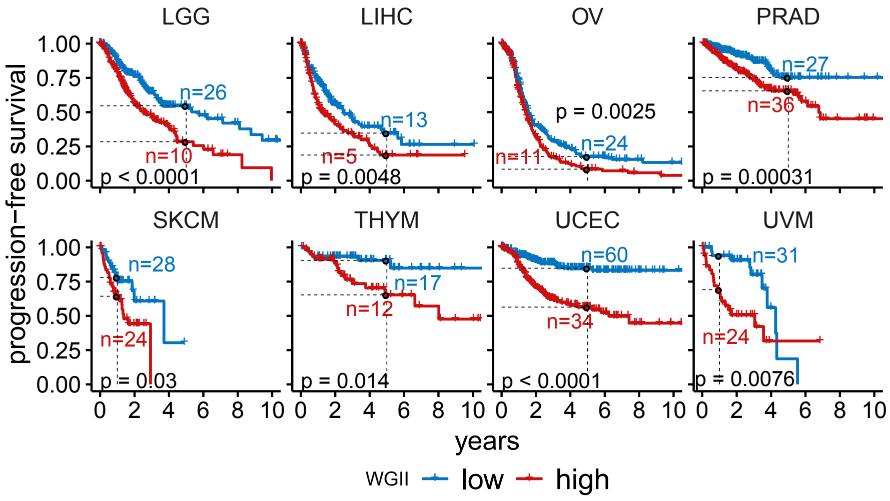


**Figure 8.** Progression free survival in eight cancer types where significant differences between high and low WGII group were observed.
